# Intrabody induced cell death by targeting the *T. brucei* cytoskeletal protein *Tb*BILBO1

**DOI:** 10.1101/2021.07.18.452872

**Authors:** Christine E. Broster Reix, Miharisoa Rijatiana Ramanantsalama, Carmelo Di Primo, Laëtitia Minder, Mélanie Bonhivers, Denis Dacheux, Derrick R. Robinson

## Abstract

*Trypanosoma brucei* belongs to a genus of protists that cause life-threatening and economically important diseases of human and animal populations in Sub-Saharan Africa*. T. brucei* cells are covered in surface glycoproteins some of which are used to escape the host immune system. Exo-/endocytotic trafficking of these and other molecules occurs *via* a single copy organelle called the flagellar pocket (FP). The FP is maintained and enclosed around the flagellum by the flagellar pocket collar (FPC). To date, the most important cytoskeletal component of the FPC is an essential, calcium-binding, polymer-forming protein called *Tb*BILBO1. In searching for novel immune-tools to study this protein, we raised nanobodies against *Tb*BILBO1. Nanobodies (Nb) that were selected according to their binding properties to *Tb*BILBO1, were tested as immunofluorescence tools, and expressed as intrabodies (INb). One of them, Nb48, proved to be the most robust nanobody and intrabody. We further demonstrate that inducible, cytoplasmic expression of INb48 was lethal to these parasites, producing abnormal phenotypes resembling those of *Tb*BILBO1 RNAi knockdown. Our results validate the feasibility of generating functional single-domain antibody derived intrabodies to target trypanosome cytoskeleton proteins.

## Introduction

Trypanosomes are flagellated protists, comprising of pathogenic species capable of infecting a wide range of vertebrate hosts including humans, domestic and wild animals. Commonly transmitted by insect vectors, trypanosomes cause lethal and economically important human and animal diseases worldwide (1, 2). A limited number of diagnostic tests and treatments are available for human and animal African trypanosomiasis and there are no vaccines. Fortunately, the first oral trypanocidal drug, Fexinidazole, was recently approved for use in humans in Africa (3, 4) which simplifies the treatment regime for certain cases. However, millions of wild animals and domesticated livestock are still at risk of infection with the economic burden of the disease to the African economy is estimated to be 4.7 billion USD per year due to the death of around 3 million head of cattle (5) (http://www.fao.org/paat/the-programme/the-disease/en/). Understanding the basic biology of any virulent organism is important to fully understand its pathogenicity. Furthermore, new targets, tools, and models are needed.

Trypanosomes possess a highly organized cytoskeleton that comprises primarily of microtubules, but also many cytoskeleton-associated proteins (5–13). This ensemble of proteins coordinates changes in cell morphology during the cell cycle, as well as permitting cell motility and reorganization during differentiation between life-cycle stages. Most pathogenic trypanosomes possess a single copy organelle called the flagellar pocket (FP), an infolding of the pellicular membrane at the site where the flagellum exits the cell body, which is physically connected to the cytoskeleton (13–18). In many species the FP is the sole site of exo- and endocytosis. In *T. brucei* the FP is essential for survival and this has been demonstrated through studies of an FP-linked cytoskeleton component called the flagellar pocket collar (FPC). The FPC is a cytoskeletal structure that circumvents the neck of the FP at the site where the flagellum exits the cell (13, 20–26). The FPC forms a cytoskeletal boundary or interface at the site of contact between the pellicular, flagellar and FP membranes (13, 20, 23–26). To date, only a handful of FPC and FPC-associated proteins have been identified and only two essential proteins have been characterized. The first is *Tb*BILBO1 (67kDa; 587aa) which is the main protein component of the flagellar pocket collar (FPC), and the second is *Tb*MORN1 (40.9kDa; 358aa) which is present in the Hook Complex (HC), a structure intimately linked and immediately distal to the FPC, involved in facilitating protein entry into the cell. Thirdly, *Tb*FPC4 (25), a microtubule binding HC protein, is a binding partner of *Tb*BILBO1 with a role in FPC segregation (14, 20, 27–30). Analysis of *Tb*BILBO1 has demonstrated its uniqueness as a FPC component and it is indeed kinetoplastid-specific. The N-terminus of *Tb*BILBO1 adopts a ubiquitin-like fold, followed by two calcium-binding EF-hand domains (EFh: amino acid residues 185–213 and 221–249). These domains are followed by a large coiled-coil domain (CC, amino acids 263-566) and a C-terminal leucine zipper (LZ, amino acid residues from 534–578) that are respectively involved in dimerization and polymerization (14, 21, 22, 26). RNA interference (RNAi)-mediated knock-down of *Tb*BILBO1 in the cultured, insect procyclic form (PCF) parasites prevents FPC and FP biogenesis, induces the aberrant repositioning of the new flagellum (detached from the cell body along its length), and results in parasite cell death. Knock-down of *Tb*BILBO1 in the mammalian bloodstream form (BSF) is rapidly lethal (14).

Antibodies are especially important tools for studying protein function in cells. A specific class of heavy chain only antibodies (HCAb) are found naturally in Camelidae (alpacas, camels, llamas and vicunas) and also in Elasmobranchs (cartilaginous fish, including sharks and rays) (31, 32). Nanobodies are recombinant antibody fragments derived from the antigen binding domain (variable domain, VHH) of heavy chain only antibodies (33, 34). They are much smaller than traditional antibodies and are ∼ 15kDa in mass (35–37). The use of nanobodies is greatly expanding in cellular and molecular biology, diagnostics, and medicine (38–44). Indeed in 2018, caplacizumab was the first nanobody-based drug, approved for treating thrombotic thrombocytopenic purpura and thrombosis (45, 46). Intrabodies (INb), nanobodies expressed within the cytoplasm of the target cell, have many potential roles ranging from pathogen detection, to treating osteoarthritis and killing of tumour cells expressing cancer specific antigens (47). Studies have shown that a single chain antibody fragment (scFv) can have knockdown effects without having a direct neutralizing effect by trapping the target membrane or secretory protein within the endoplasmic reticulum (48). By using targeting motifs, intrabodies can also be directed to specific organelles or structures.

Intrabodies have the benefit of having high antigen specificity, the ability to target a specific cell compartment, a particular isoform or post-translational modification, the ability to recognize and bind to a structural epitope not accessible by conventional antibodies, and have potential as tools for many biological systems (49, 49–53). These attributes have increased the potential for therapeutic intrabodies for neurological disease, cancer (VEGF) viral infections (HIV) and autoimmune disease (TLR9) (39, 42, 54–58). At the time of writing, there are currently thirteen single domain antibodies (from both camelid and shark origin) in either clinical or preclinical trials for treatment of a variety of conditions such as rheumatoid arthritis, solid tumors and respiratory disease (https://clinicaltrials.gov). Numerous important studies have used nanobodies to characterize trypanosome proteins as well as anti-trypanosome drug or toxin carriers, whilst others have proved to have intrinsic lytic activity (40, 43, 53, 59–69). To date, only one nanobody has been developed against a trypanosome cytoskeleton protein, Nb392, targeting the paraflagellar rod protein (PFR1), a structure that runs along the axoneme within the flagellum. Nb392 proved to be an excellent tool for immunofluorescence, protein isolation and a potential diagnostic tool and demonstrating the versatility of Nb against parasite cytoskeletal proteins (66).

The FPC is intimately associated with the MtQ; a subset of specialized microtubules that originate near the basal bodies of the flagellum and grow very close to, if not through, the FPC (71, 72). Previous published work has identified binding of *Tb*MORN1 and *Tb*SPEF1 (a microtubule binding and bundling protein) onto detergent and salt-extracted MtQ (20, 25, 73). Prior to those studies, our anti-*Tb*BILBO1 (IgM) antibody demonstrated both an FPC and a minor MtQ signal. We show here that, our anti-*Tb*BILBO1 (1–110) rabbit polyclonal, purified Nb48 and internally expressed nanobody 48 (INb48) all bind to the FPC and the MtQ. Importantly, we demonstrate that nanobodies (intrabodies) raised against *Tb*BILBO1 can function as potential knock-down agents and we demonstrate that anti-*Tb*BILBO1 intrabody (INb48) identifies its target protein when expressed in an homologous mammalian system *in vitro* and within the cytoplasm of *T. brucei.* Furthermore, INb48 prevented FPC formation and function and induced parasite cell death.

## Results

### Production of anti-*Tb*BILBO1 nanobodies

Full-length histidine-tagged *Tb*BILBO1 protein was expressed in bacteria. Purified recombinant protein was then used to immunize an alpaca to produce *Tb*BILBO1 targeting nanobodies by the Nanobody Service Facility (NSF, VIB, Brussels). Seven different Nbs were identified following phage display screening. These were assigned to three different groups based on their amino acid sequences. It was hypothesized that Nbs in the same group would recognize the same epitope, but their other characteristics such as affinity, potency, stability and expression yield, could be different. We chose to focus on one nanobody from each group for our current studies: Nb48, Nb9, and Nb73.

### Nb48 as a functional tool for immunolabelling *Tb*BILBO1

The Nbs were cloned in the pHENC6c vector in frame with a HA and Histidine tag (Nb_::HA::6His_) allowing the secretion of the Nbs in the periplasm, their purification using metal affinity chromatography, and detection. The Nbs were first tested by western blotting (Figure 1A). Samples of non-induced (-) and induced bacteria (+), pellet (P) and periplasmic extracts (Ex) were probed by western blot with anti-HA (Figure 1A and Supplementary information S1A). The periplasmic extracts, containing the soluble anti-*Tb*BILBO1 Nbs, were purified using metal affinity chromatography and these nanobodies were probed by western blot. As an example, samples from the purification steps of Nb48_::HA::6His_ probed by western blot with anti-Histidine (anti-His) are shown in Figure 1B. For simplicity, the HA::6HIS tagged nanobodies are hereafter called Nb48, Nb9 and Nb73 (with tags written in subscript). We tested purified Nb48_::HA::6His_, Nb9_::HA::6His_ and Nb73_::HA::6His_ by using anti-HA for detection in immunofluores-cence microscopy (IFA) on *T. brucei* wild-type (WT) procyclics (PCF) detergent-extracted cells (cytoskeletons) (Figure 1C-F). For convenience, in Figure 1 and 2, the cells that are probed with purified nanobodies are labelled as “Anti-Nb”, plus the relevant nanobody name. In these experiments purified Nb48_::HA::6His_ labels endogenous *Tb*BILBO1 at the FPC, as seen by co-localization with anti-*Tb*BILBO1 (polyclonal antibody 1-110) (Figure 1D). Nb9_::HA::6His_ does not label wild-type cytoskeletons expressing endogenous *Tb*BILBO1 protein (Figure 1D) but does label *Tb*BILBO1 in cells that over-express *Tb*BILBO1::3cMyc (Figure 1E). Importantly, Nb73_::HA::6His_ was negative in our experiments and did not label *Tb*BILBO1 in either endogenous (Figure 1F) or overexpressed levels of *Tb*BILBO1 (not shown). To exclude the possibility of cross-reaction between primary and or secondary antibodies in double labelling protocols, Nb48_::HA::6His_ and anti-*Tb*BILBO1 (1–110) were also used alone to probe cytoskeletons. In both cases the subsequent labelling was observed on the FPC and the microtubule quartet, (MtQ) a subset of specialized microtubules that are in close proximity to the FPC (71, 72), (Figures 2A and B). Labelling with anti-*Tb*BILBO1 (rabbit polyclonal, 1-110) alone also clearly labelled the MtQ (Figure 2C). Co-localization was observed in double labelling experiments with anti-*Tb*BILBO1 (1–110) and Nb48_::HA::6His_ (Figure 2D). In Figures 2A-D, the asterisks indicate the labelling of the MtQ. Purified Nb48_::HA::6His_ positively labels full-length *Tb*BILBO1 protein from *T. brucei* whole cell extracts in western blotting experiments (Figure 2E) as well as *Tb*BILBO1 truncated versions (T3::3cMyc, aa 171-587 and T4::3cMyc, aa 251-587) (please also refer to the schematic in Figure 2F for an overview of the truncations). Nb48_::HA::6His_ did not label T1 or T2 (not shown) indicating that Nb48 binds to *Tb*BILBO1 in the region ranging from aa 251 to aa 587. Controls for IFA experiments in Figures in 1 and 2 are presented in Supplementary information S1B.

**Fig 1.**
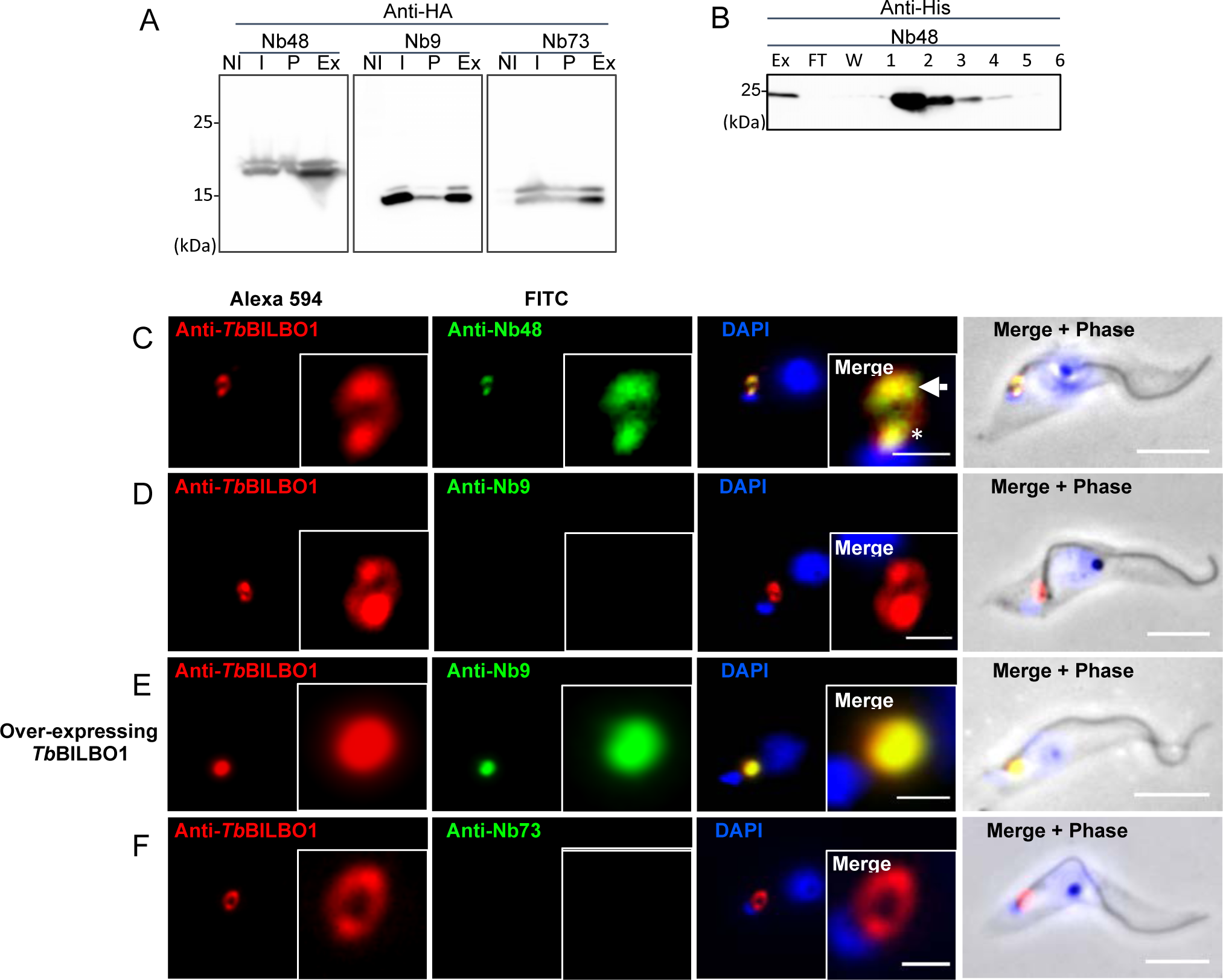
Anti-*Tb*BILBO1 nanobodies: purification and use in IFA. (A) Bacterial expressed nanobody (Nb_::HA::6His_) samples probed on a WB with anti-HA. Samples of non-induced (NI), induced (I) bacterial cultures, pellet (P) and periplasmic extract (Ex). (B) Purification of Nb48_::HA::6His_ : flow through (FT), wash (W) and elutions (lanes 1-6), all lanes were probed by WB using anti-His. (C) IFA testing of purified Nb48_::HA::6His_ (green) on wild-type (WT) *T. brucei* cytoskeletons showing co-localization with *Tb*BILBO1 (probed with anti-*Tb*BILBO1 aa1-110 rabbit; red). The arrow indicates the FPC and the asterisk indicates co-labelling on the MTQ; both of these structures are labelled by Nb48_::HA::6His_ and anti-*Tb*BILBO1 1-110 rabbit. (D) Purified Nb9_::HA::6His_ does not label endogenous *Tb*BILBO1 in WT cytoskeletons. (E) Nb9_::HA::6His_ on *T. brucei* over-expressing *Tb*BILBO1 (++) shows co-localization with *Tb*BILBO1. (F) Purified Nb73_::HA::6His_ on WT cytoskeletons does not label *Tb*BILBO1. Scale bar = 5μm, inset = 1μm. For the sake of convenience the protein being probed is named as anti-Nb, plus the relevant nanobody name rather than its tag.

**Fig 2.**
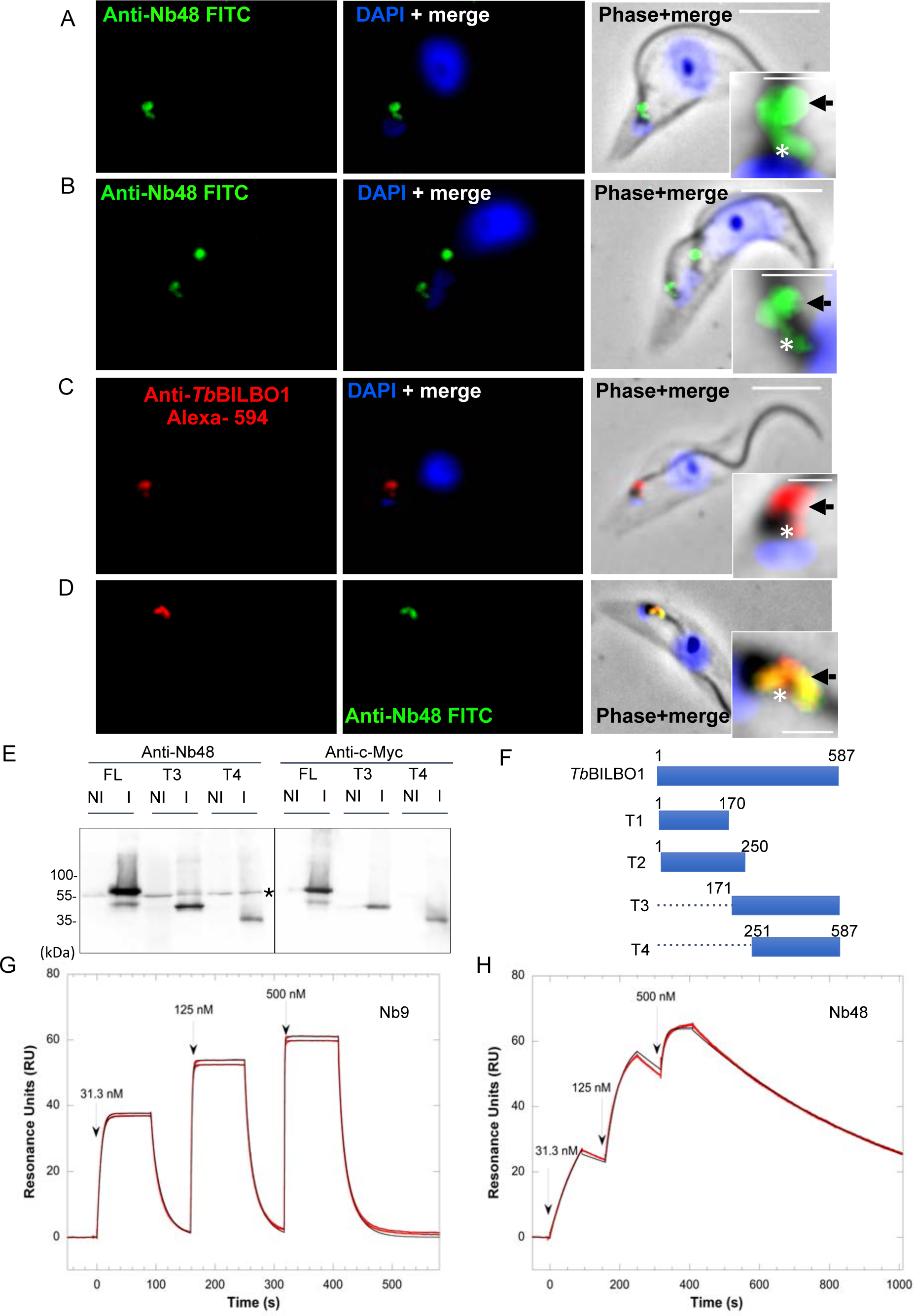
Anti-*Tb*BILBO1 nanobodies Nb48_::HA::6His_ and Nb9_::HA::6His_ bind to *Tb*BILBO1 *in vitro*. (A) and (B) show WT PCF cytoskeletons (94), probed with Nb48_::HA::6His_ followed by anti-HA, then FITC conjugated anti-mouse. (A) is a parasite cell in the 1K1N stage of the cell cycle (K=kinetoplast, n=nucleus). (B) is a cell in the 2K1N stage. Both cell stages show Nb48_::HA::6His_ labelling on the FPC (black arrow) and MtQ (white asterisk). The MtQ signal was not always apparent on every cell imaged, which could be due to cell cycle stage. (C) WT *T. brucei* cytoskeleton probed with anti*-Tb*BILBO1 (1–110) followed by anti-rabbit Alexa fluor 594. (D) WT cytoskeleton probed with anti-*Tb*BILBO1 and Nb48_::HA::6His_ showing complete co-localization. (E) Western blot analysis of whole cell extracts of *T. brucei* PCF cells expressing *Tb*BILBO1 full-length (FL) and *Tb*BILBO1 truncations, probed with Nb48_::HA::6His_. Non-induced (NI) and induced (I) trypanosomes expressing *Tb*BILBO1::3cMyc FL protein, aa 171-587 (T3), and aa 251-587 (T4) were probed with purified Nb48_::HA::6His_. Note: all proteins are running slightly faster than predicted size; FL should run at 70kDa, T3 should run at 49kDa and T4 at 40kDa. The black asterisk indicates labeling of endogenous *Tb*BILBO1 seen across all samples. (F) A schematic diagram of cMyc tagged *Tb*BILBO1 full-length (FL) and truncations. T1 and T2 are soluble. Nb48_::HA::6His_ binds to a region within aa 251 and 587 of *Tb*BILBO1. (G) Kinetic analysis by surface plasmon resonance (SPR) of Nb9_::HA::6His_ and (H) Nb48_::HA::6His_, binding to 6His::*Tb*BILBO1. The results of Nb9_::HA::6His_ and Nb48_::HA::6His_ binding are shown on the sensorgrams respectively, and are represented by the red curves. The black lines represent the theoretical fit of each Nb as obtained from the Biaevaluation software to a kinetic titration data set of three concentrations of the nanobodies *in vitro.* Nb9_::HA::6His_ displays a good affinity for 6His::*Tb*BILBO1 with a K**D** of 15.2 ± 0.7 nM, however, Nb48_::HA::6His_ displays the highest affinity for 6His::*Tb*BILBO1 with a KD of 8.8 ± 0.7 nM. For the sake of convenience the protein being probed is named as anti-Nb, plus the relevant nanobody name rather than its tag. Scale bar = 5μm, inset = 1μm.

### Surface Plasmon Resonance confirms a strong binding affinity of Nb48 to *Tb*BILBO1

To determine the equilibrium dissociation constant (KD) of Nb48_::HA::6His_ (group 1) and Nb9_::HA::6His_ (group 2) to *Tb*BILBO1, we used surface plasmon resonance (SPR), (Figures 2G and H), (note: It was not possible to assess Nb73_::HA::6His_ (group 3) using SPR due to the inconsistent yield and impure production of this nanobody from bacteria). Sensor chips were coated with purified 6xHis::*Tb*BILBO1 by amine coupling and probed sequentially at increasing concentrations with purified Nb9_::HA::6His_ and Nb48_::HA::6His_ The Nb binding results are shown on the sensorgrams (Figures 2G and H) respectively, and are from a titration data set of three concentrations of the nanobodies: 31.3nM, 125nM and 500nM. These results illustrate that Nb9_::HA::6His_ and Nb48_::HA::6His_ bind to 6xHis::*Tb*BILBO1, but do not behave kinetically in the same way. The superposition of the sensorgrams with the fitting curves validates the Langmuir 1:1 model of interaction for analyzing them. Nb9_::HA::6His_ displays faster rate constants than Nb48_::HA::6His_ with Nb9_::HA::6His_ *k_a_*=6.54 x 10^6^ ± 0.27 x 10^6^ M^-1^ s^-1^ and *k_b_*=98.9 x 10^-3^ ± 0.09 x 10^-3^ s^-1^, and Nb48_::HA::6His_ *k_a_*=1.74 x 10^5^ ± 0.16 x 10^5^ M^-1^ s^-1^ and *k_b_*=1.52 x 10^-3^ ± 0.02 x 10^-3^ s^-1^. Because of a 100x fold slower rate of dissociation, despite a 10x fold slower rate of association, Nb48_::HA::6His_ displays the highest affinity for 6xHis::*Tb*BILBO1 with a K_D_ of 8.8 ± 0.7 nM compared to Nb9_::HA::6His_ K_D_ of 15.2 ± 0.7 nM. These values confirm that Nb48_::HA::6His_ has the highest affinity for 6xHis::*Tb*BILBO1. Nb48 provided the best results as a tool by IFA, as a probe on western blot and binding to *Tb*BILBO1 as measured by SPR, therefore we decided to focus on Nb48 for further studies.

### Anti-*Tb*BILBO1 intrabodies bind to the FPC and the MtQ

Previous studies using STED and immuno-electron microscopy illustrated that the anti-*Tb*BILBO1 (1–110) rabbit polyclonal antibody showed labelling both at the FPC and along the MtQ (25). Given that *Tb*BILBO1 and *Tb*MORN1 are stable on isolated flagella, we questioned whether the FPC could be probed i*n vivo* in trypanosomes by intra-nanobody (intrabody, INb) targeting. The DNA sequence encoding Nb48 plus a C-terminal 3cMyc tag was cloned into a *T. brucei* expression vector allowing the tetracycline-inducible expression of the recombinant Intrabody48 (INb48::3cMyc). Isolated flagella derived from procyclic cells (PCF) expressing INb48::3cMyc for 12 hours were prepared for IFA and double labelled by probing with anti-cMyc and anti-*Tb*BILBO1 (1–110), showing co-localization of *Tb*BILBO1 and INb48 (Figure 3A). For convenience the use of INb48 is displayed in these figures as Anti-INB48 rather than indicating the tag used on the intrabody. Immuno-electron microscopy was also carried out flagella that were double labelled using anti-cMyc and anti-*Tb*BILBO1 antibodies (Figure 3B). Both the IFA and the electron micrographs of these experiments revealed that INb48::3cMyc colocalizes with anti-*Tb*BILBO1 on the FPC, but also on the proximal region of the MtQ as described earlier in the IFA studies shown in Figures 2A-D. Of note, Gheiratmand *et al.,* 2013, Esson *et al.,* 2012 and Albisetti 2017 *et al.,* have shown that *Tb*SPEF1 (a microtubule binding and bundling protein) and *Tb*MORN1, are localized to the MtQ of flagella thus clearly indicating that microtubule binding proteins do remain attached to this structure after harsh extraction (73, 20, 25). We also carried out temporally shorter expression assay of INb48::3cMyc which revealed that it is observed on the FPC and the MtQ at 3 hours post induction (Supplementary information SI Figure 1B). No first antibody controls are also shown in supplementary information - SI Figure 1C).

**Fig 3.**
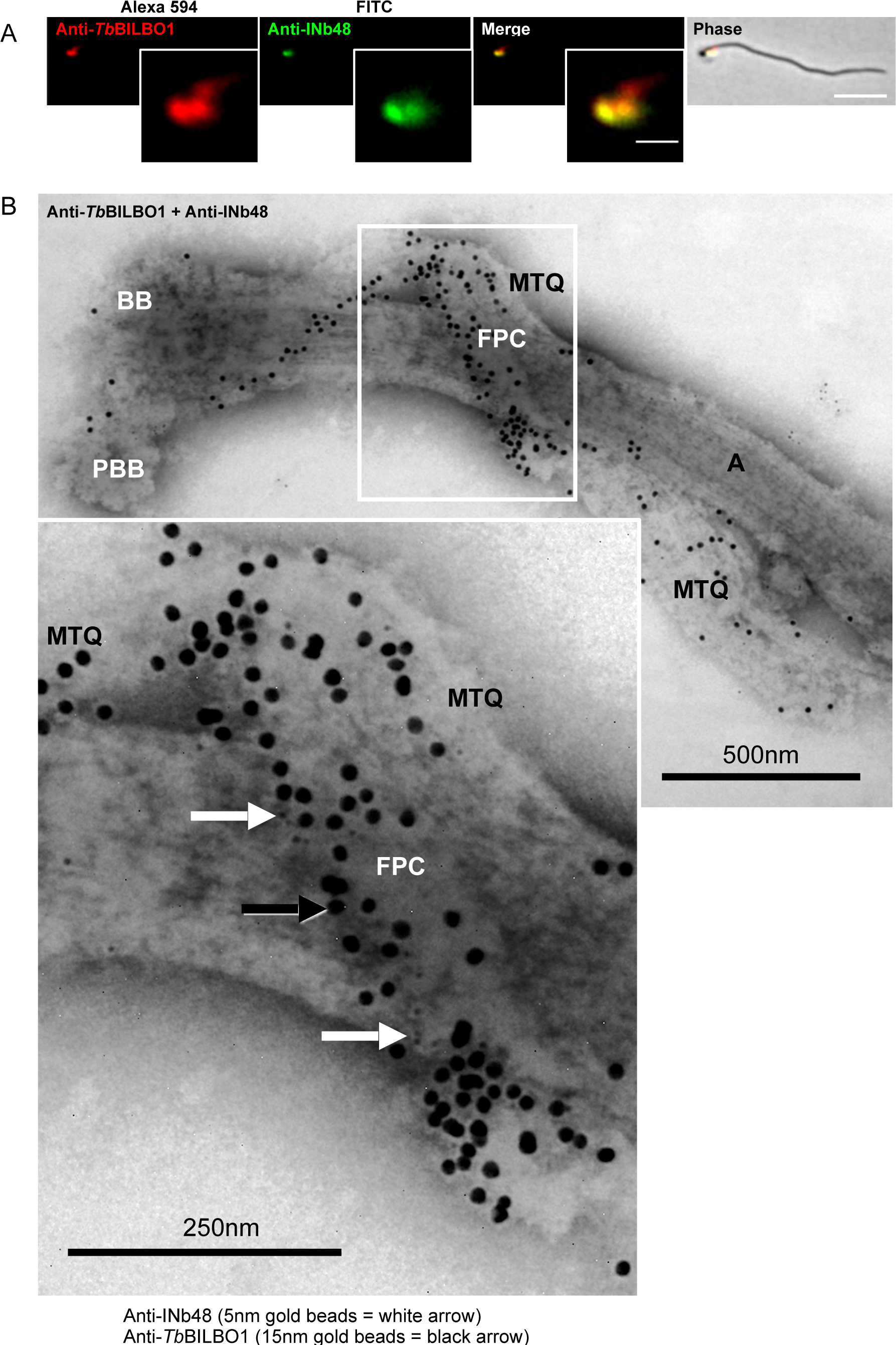
INb48::3cMyc expression targets the FPC in *T. brucei*. (A) Co-localization of INb48::3cMyc with *Tb*BILBO1 on isolated *T. brucei* PCF flagella. Scale bar = 5μm, inset = 1μm. (B) Immuno-electron micrographs of a purified flagellum from a trypanosome expressing INb48::3cMyc3 for 24hpi showing clear co-localization of INb48::3cMyc (5nm gold beads = white arrow) with *Tb*BILBO1 (15nm gold beads = black arrow). Both 5nm and 15nm sized gold beads are also clearly visible on the MtQ. BB = Basal Body, PBB = Pro-Basal body, FPC = Flagellar Pocket Collar, MtQ = Microtubule Quartet, A= axoneme. Scale bar = 500nm, inset = 250 nm. For the sake of convenience the protein being probed is named as anti-Nb, plus the relevant nanobody name rather than its tag.

### The expression of anti-*Tb*BILBO1 intra-nanobodies is cytotoxic in *T. brucei*

Because anti-*Tb*BILBO1 intrabodies co-localize with *Tb*BILBO1 on isolated flagella, we hypothesized that INbs bind tightly to *Tb*BILBO1 and, as a consequence, may disturb FPC formation or function. To test this hypothesis, we monitored the effect of expression of INb48::3cMyc in PCF *T. brucei* using different concentrations of tetracycline (Figure 4A). Tetracycline concentrations of 0.1 ng/mL and 1ng/mL had no impact on cell growth when compared to the non-induced cells (-Tet; red line). However, concentrations from 5ng/mL to 1*μ*g/mL induced a rapid growth defect from 24 hours post induction (hpi), and cell death after 24hpi. The change in growth rates related to different doses of tetracycline indicates that the lethality of INb48::3cMyc expression in PCF is dose dependent. Taking in account these results, we chose a tetracycline concentration of 1*μ*g/mL as the standard concentration for INb48::3cMyc, INb9::3cMyc and INb73::3cMyc induction. After induction of the expression of these three INbs in trypanosomes, daily samples were taken for cell counts (Figure 4B-D) and western blot analysis (Figure 4E). Independent induction of each INb resulted in phenotypes that ranged from rapid cell death from 24hpi for INb48::3cMyc,(Figure 4B), to a reduction in cell growth rate for INb9::3cMyc (Figure 4C) or no change in growth rate for INb73::3cMyc (Figure 4D). Western blot analysis of INb expression is shown in (Figure 4E) and illustrates robust expression of all INbs at 24 hours post induction in trypanosomes. After 48hpi of intrabody expression, the signals for INb48::3cMycand INb9::3cMyc were lost and the signal for *Tb*BILBO1 and tubulin control proteins show signs of degradation, indicating some degree of cell death in the population. However, little or no degradation is observed in cells clearly expressing INb73::3cMyc which, importantly, showed no effect on cell growth.

**Fig 4.**
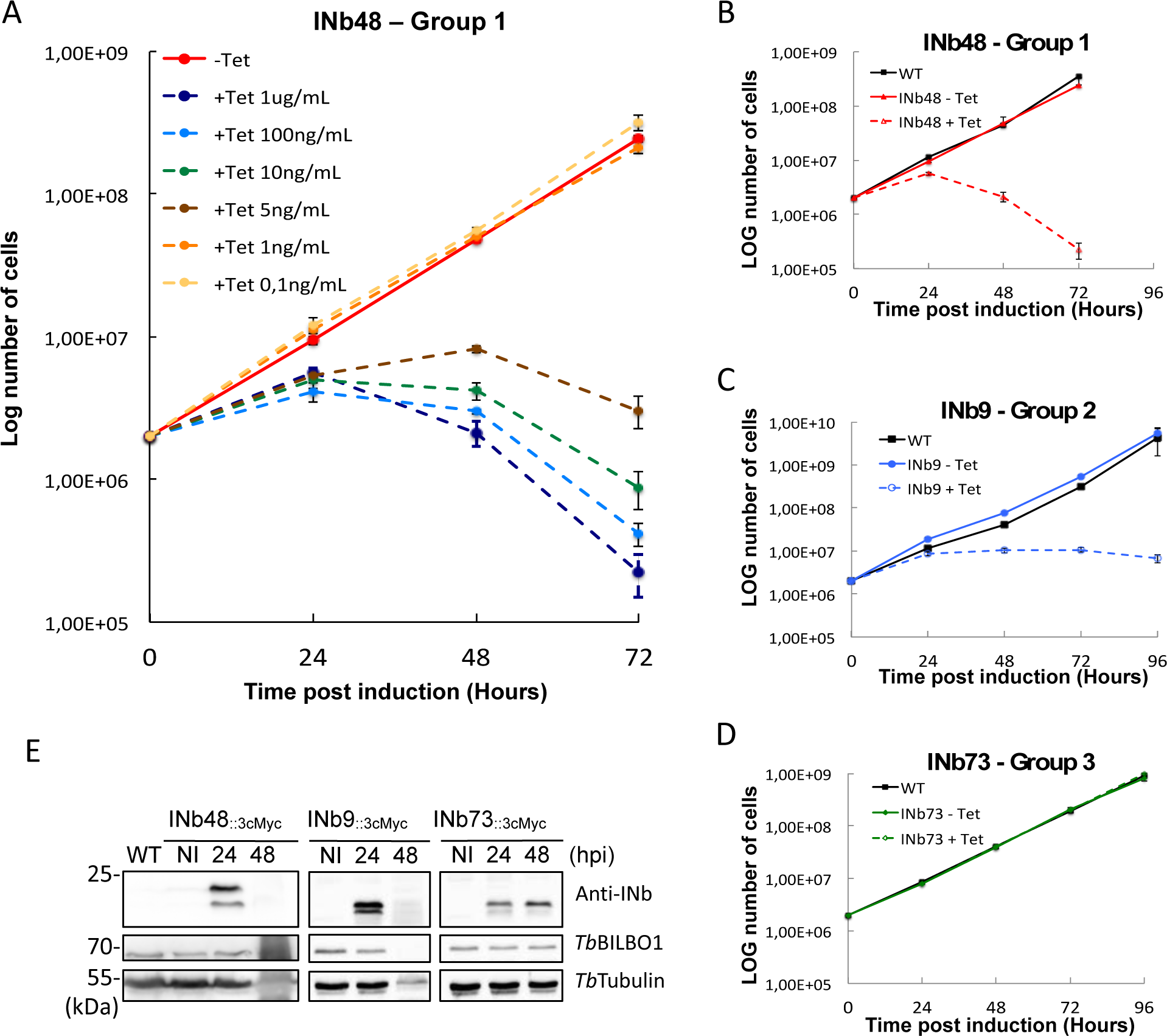
INb48::3cMyc expression is trypanocidal. (A) Growth curves of trypanosome cells expressing INb48::3cMyc. Tetracycline dose dependent effect on trypanosome growth and cytotoxicity for INb48::3cMyc expression in *T. brucei.* (B) INb48::3cMyc (group 1) expressed in PCF induces rapid decline in growth and trypanosome cell death, (C) INb9::3cMyc (group 2) expressed in PCF induces cell growth arrest and cell death, and (D) INb73::3cMyc (group 3) expressed in PCF has no effect on cell growth. (E) Western blot of INb48::3cMyc, INb9::3cMyc and INb73::3cMyc expression in PCF, showing expression only in induced cells. Rapid degradation of protein due to cell death at 48hpi is observed for INb48::3cMyc and INb9::3cMyc. Time post induction is in hours.

As a negative control an intrabody against Green Fluorescent Protein (INbGFP::3cMyc) was expressed in trypanosomes, and as expected, no cell growth defect or phenotype was observed despite its strong expression and cytoplasmic localization (Supplementary Figure 2A-C). These data demonstrate that expression of INb48::3cMyc and of INb9::3cMyc are lethal, with INb48::3cMyc showing the strongest effect, whilst expression of INb73::3cMyc had no effect on cell growth. These data suggest that INb48::3cMyc and INb9::3cMyc bind to *Tb*BILBO1 *in vivo,* and hinder its function.

### Intrabody 48 (INb48::3cMyc) binds to *Tb*BILBO1 and induces *Tb*BILBO1 RNAi knockdown phenotypes

Trypanosome cell cycle stages can be defined based on the arrangement and number of nuclei and mitochondrial genome (kinetoplasts) present in the cell. In wild-type G1 PCF cells, the linear organization of kinetoplast (K), and the nucleus (N) is: kinetoplast followed by the nucleus (1K1N). As the cell passes through the cell cycle the kinetoplast duplicates to form 2K1N cells, then mitosis and nuclear duplication ensues to form 2K2N cells with a posterior new flagellum (NF) and a more distal old flagellum (OF), followed by cytokinesis producing 1K1N mother and daughter cells. This means that wild-type cells should never possess more than the 2K2N DNA complement (73). RNAi knockdown of *Tb*BILBO1 in cultured PCF cells results in cell-cycle arrest at the 2K2N stage, producing cells with elongated posterior ends, new flagella that were detached from the length of the cell body, and inhibition of new FP formation (13). We investigated further the localization of INb48 and its effects on cell morphology when expressed in *T. brucei,* (Figure 5). Figure 5A shows non-induced (- Tet, NI), whereby the NI trypanosomes display the classic immunofluorescence signal for *Tb*BILBO1 with the annular labelling at the FPC and along the MtQ between the FPC and the basal bodies (BB). After 24hpi of INb48::3cMyc (+ 24h) cells, unusual morphological phenotypes were observed whereby the new flagellum was detached from the length of the cell body and cells demonstrated elongated posterior ends. After expression of INb48::3cMyc, both INb48::3cMyc and *Tb*BILBO1 were observed at the old FPC and sometimes also at the base of the detached new flagellum (Figure 5B, asterisk). Rather than the classic annular and MtQ signals observed after anti-*Tb*BILBO1 labelling, we observed both *Tb*BILBO1and INb48::3cMyc signals extending along the cell (Figure 5B, merged inset). This result demonstrates that no new FPC was formed after INb48::3cMyc expression and that INb48::3cMyc binding to *Tb*BILBO1 disturbs FPC formation.

**Fig 5.**
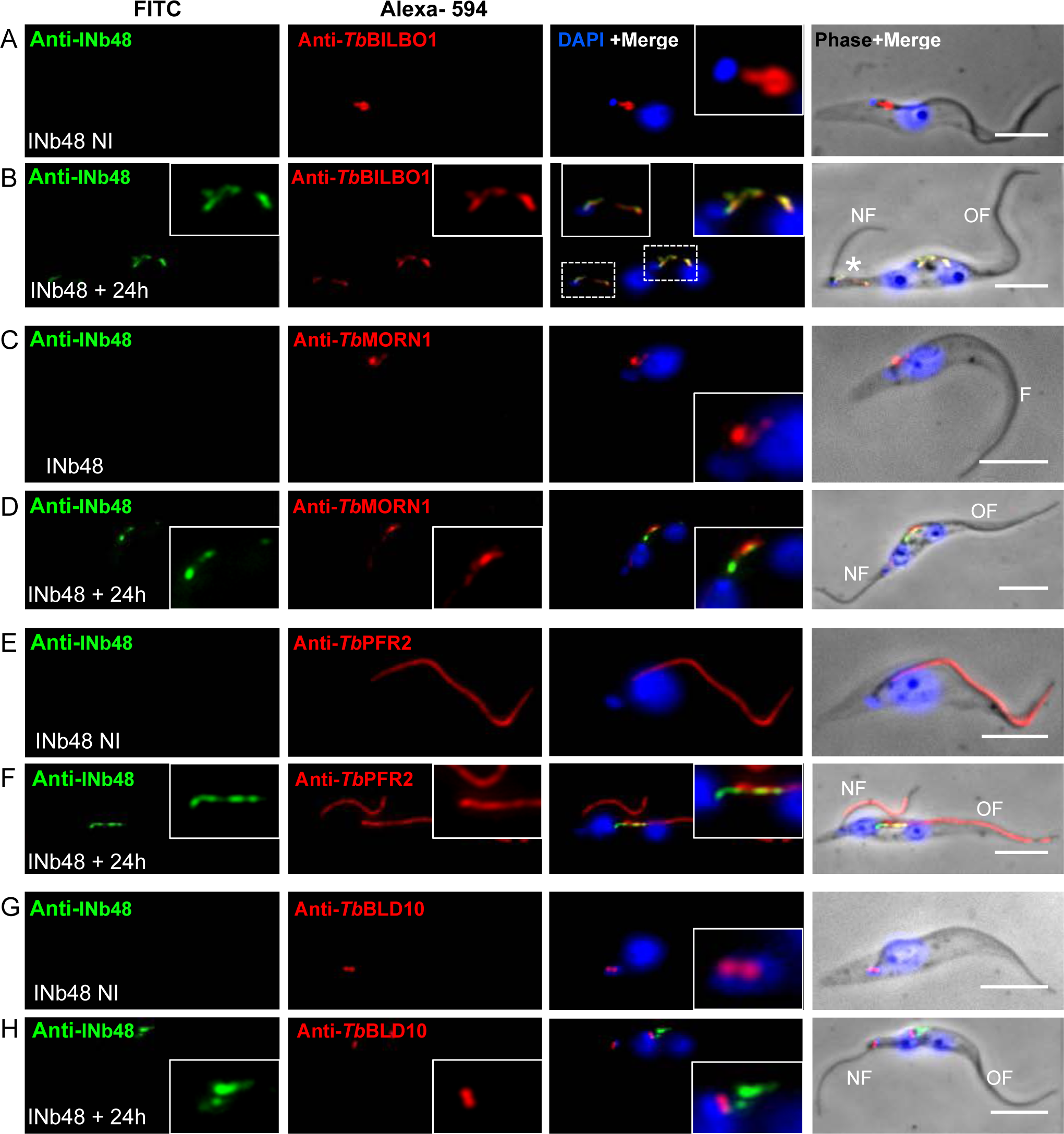
INb48**::3cMyc** expression *in vivo* induces phenotypes resembling *Tb*BILBO1 RNAi knockdown. (A) IFA of non-induced (NI) PCF *T. brucei* cells probed with anti-cMyc as a probe for INb48::3cMyc (green) and anti-*Tb*BILBO1 (red), indicating the absence of INb48::3cMyc labelling in non-induced cells. (B) An INb48::3cMyc induced (Ind) *T. brucei* cell at 24hpi showing co-localisation of INb48::3cMyc (green) with a disorganized signal for *Tb*BILBO1 (linear polymer formation instead of the typical annular shape) and a posteriorly localised, detached new flagellum (NF). Asterisks (*) denote the presence of co-localisation of INb48::3cMyc and anti-*Tb*BILBO1 is observed on the old flagellum (OF) and at the base of the new detached flagellum (NF). (C) Non-induced NI) trypanosome showing correct location of *Tb*MORN1 labelling (red) at the base of the flagellum (F), with the classic hook and shank composition; there is no anti-cMyc signal in this NI cell. (D) At 24hpi (Ind), INb48::3cMyc labelling (green) is seen in the location of the FPC with linear labeling extending a short way along the axoneme; *Tb*MORN1 labelling has lost the classic hook shape and is seen as an extended linear form extending distally from the FPC region along the direction of the old flagellum (OF). (E) Non-induced (NI) trypanosome showing typical labelling of *Tb*PFR2 (red) along the flagellum. (F) 24hpi Ind INb48::3cMyc (green) showing a trypanosome cell with both the old (OF) and new flagella (NF) labelled with *Tb*PFR2. INb48::3cMyc labelling is observed only at the old FPC extending along anteriorly in this cell. (G) A non-induced (NI) trypanosome cell probed with anti-*Tb*BLD10 (red), labelling both the mature and pro-basal bodies (BB) at the base of the flagellum. (H) At 24hpi (+Tet) of INb48::3cMyc (green), a detached new flagellum (NF) is seen the tip of at an extended posterior end; anti-*Tb*BLD10 labelling is seen associated with both the new and old flagella (red); INb48::3cMyc labelling is seen mainly in the region of the FPC of the old flagellum (green). Scale bar = 5μm. For the sake of convenience the protein being probed is named as anti-Nb, plus the relevant nanobody name rather than its tag.

Because FP biogenesis is hindered in INb48::3cMyc induced cells, we decided to investigate the organization of the hook complex (HC), a structure intimately associated with the FPC, by immunolabelling of *Tb*MORN1 (28) in the background of INb48::3cMyc induced mutant cells. The HC is an essential structure positioned distal to the FPC, forming a distinctive hook shape, and is involved in the regulation of entry of molecules into the flagellar pocket (20, 27, 28, 30). Figure 5C shows the normal location and hook-like shape of anti-*Tb*MORN1 labelling in non-induced cells. However, at 24hpi of INb48::3cMyc expression, *Tb*MORN1 labelling is elongated distally along the flagellum, and the hook shape is lost (Figure 5D). Importantly, little or weak *Tb*MORN1 labelling is observed at the base of the new flagellum, suggesting that HC biogenesis is also strongly affected in cells expressing INb48::3cMyc,. A consistent observation in cells expressing INb48::3cMyc was that the new flagellum was detached from the length of the cell body, *i.e.* only attached at the base, an abnormal phenotype strongly resembling the *Tb*BILBO1 RNAi knockdown phenotype (14). We characterized this detached flagellum phenotype by co-labelling with anti-*Tb*PFR2 antibody (8). *Tb*PFR2 is a protein of the paraflagellar rod (PFR) that is present alongside the axoneme of the flagellum of wild-type *T. brucei* (74) as shown in non-induced cells (Fig 5E). In induced cells (24hpi), *Tb*PFR2 labelling was observed along the old flagellum and the newly detached new flagellum (Figure 5F). In these cells, INb48::3cMyc was only observed at the old FPC, indicating that a new flagellum and PFR was formed in the absence of the FPC. To investigate any effect of INb48::3cMyc expression on the basal bodies (BB), an antibody marker anti-*Tb*BLD10 was used, which was raised against *Tb*BLD10, an essential protein in pro-basal body biogenesis (75). Figure 5G shows a non-induced *T. brucei* cell with the typical labelling of the mature and pro-basal bodies. At 24hpi of INb48::3cMyc, the labelling of the both the pro- and mature basal bodies of the mother and daughter flagella were unaffected (Figure 5H).

The phenotypes induced after INb48::3cMyc expression in trypanosomes were observed in more detail by thin section transmission electron microscopy (TEM) (Figure 6A-C). TEM micrographs of thin sections through the FP (*) of a wild-type cell (Figure 6A) illustrate that the flagellum passes through this structure and that its transition zone is positioned within the FP lumen (black arrow). INb48::3cMyc expression for 24 hours revealed that many of the new flagella were formed in the absence of an FP (Figure 6B, black arrowheads indicate the position where the FP should be formed), and the transition zone is now external to the cell (black arrows). These cells also accumulated cytoplasmic vesicles (Figure 6C). These abnormal features are typical of *Tb*BILBO1 RNAi knockdown phenotypes (14) indicating that *Tb*BILBO1 function was indeed hampered by the expression of INb48::3cMyc. We isolated flagella from INb48::3cMyc trypanosome cells at 24hpi (+Tet) and probed them with anti-cMyc (for INb48::3cMyc) and anti-*Tb*BILBO1 or, in a separate experiment, anti-cMyc and anti-*Tb*MORN1. The flagella were prepared for negative staining and visualized by transmission electron microscopy (TEM). The co-labelling of INb48::3cMyc with *Tb*BILBO1 or *Tb*MORN1 revealed that these latter proteins co-localize on the MtQ (Figure 6D and E). *Tb*BILBO1 was located on the MtQ proximal to the basal bodies (Figure 6D), whereas *Tb*MORN1 was primarily on the MtQ distal to the basal bodies (Figures 6E); *Tb*MORN1 was previously shown on the MtQ by Esson *et al.,* (20) and Albisetti *et al*., 2017 (25). Interestingly, the typical annular FPC structure is absent in INb48::3cMyc induced trypanosomes, explaining the detached flagellum phenotype. In Figure 6D, INb48::3cMyc (arrowheads) is present alongside *Tb*BILBO1 (arrows) extending from the basal bodies on the MtQ to the location where the FPC should be present. Figure 6E shows *Tb*MORN1 labelling and shows in more detail that, in the presence of INb48::3cMyc (arrowheads), it is observed on the MtQ (arrows) extending in a linear fashion proximally along the flagellum but not distally. These data demonstrate that the FPC and HC structures are perturbed by the expression of INb48::3cMyc in *T. brucei*.

**Fig 6.**
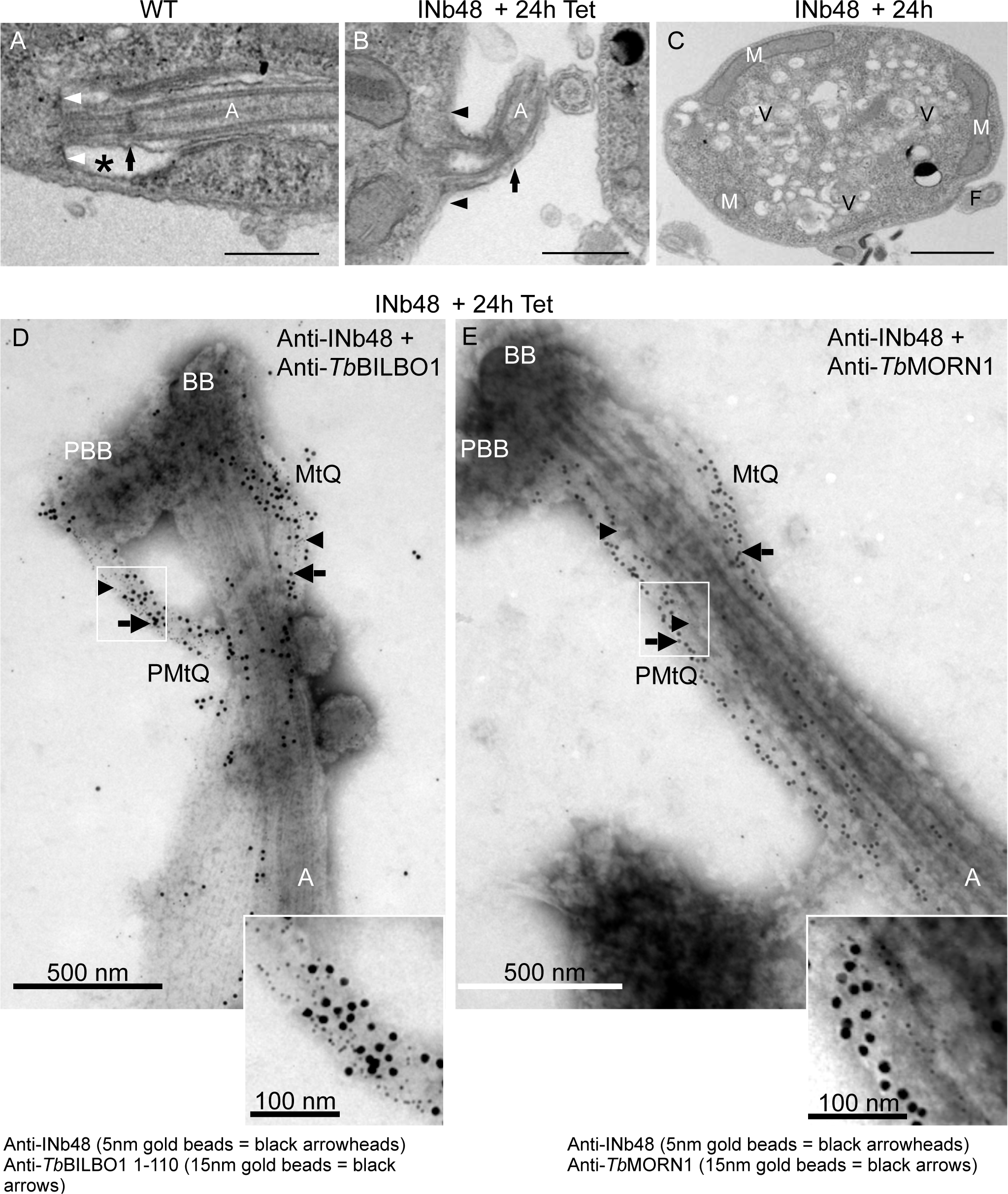
INb48**::3cMyc** expression disrupts FPC and HC formation. (A) Transmission electron micrograph (TEM) of a thin section of a WT PCF *T. brucei* cell showing the typical flagellar pocket (FP) (*), the base of the flagellar pocket (white arrows) , flagellar axoneme (A), and exit site of the axoneme from the FP membrane; the transition zone is indicated by a black arrow within the FP. (B) Illustrates a detached flagellum from a thin section of PCF cells expressing INb48::3cMyc 24hpi (+Tet). The FP has not been formed and microtubules are present where the FP membrane should be located (black arrowheads). The transition zone of the axoneme is outside the cell body (black arrow). (C) TEM of a thin section of a cell expressing INb48::3cMyc for 24hpi (+Tet), showing accumulation of intracellular vesicles. M = Mitochondrion, V = vesicles, F = flagellum. Scale bar = 1µ m. (D) Immuno-electron micrograph (IEM) of isolated flagella of PCF *T. brucei* after 24hpi (+Tet) of INb48::3cMyc expression. INb48::3cMyc is observed (5nm gold beads = black arrowheads) co-localising with *Tb*BILBO1 (15nm gold beads = black arrows) on the MtQ between the BB and the expected location of the FPC; the FPC is absent. (E) IEM of isolated flagella of PCF *T. brucei* after 24hpi (+Tet) of INb48::3cMyc expression showing INb48::3cMyc (5nm gold beads = black arrowheads) on the MtQ and an abnormal elongated labelling of *Tb*MORN1 (15nm gold beads = black arrows). Note that in trypanosome cells expressing INb48::3cMyc, the new FPC is absent and the Hook Complex as represented by *Tb*MORN1 has lost its hook-shape. BB = Basal Body, PBB = Pro-Basal body, MtQ = Microtubule Quartet, A= axoneme, PMtQ = Presumed Pro-basal body MtQ. Scale bar = 500nm, inset scale bar = 100nm. For the sake of convenience the protein being probed is named as anti-Nb, plus the relevant nanobody name rather than its tag.

Expression of INb48::3cMyc disrupts the cell cycle in PCF *T. brucei*

The linear organization of kinetoplasts and nuclei in INb48::3cMyc expressing cells and morphological phenotypes were analyzed at different time-points (Figure 7A). The percentage of non-induced (-Tet) cell cycle stages were 74.4% in the 1K1N stage (blue bars), 13.9% in the 2K1N stage, and 9.4% in the 2K2N as previously published (14). A change was observed in trypanosomes expressing INb48::3cMyc for 12 hours (+Tet 12h), with a reduction in 1K1N cells and an increase in 2K1N and 2K2N cells. There was also the appearance of abnormal K/N phenotypes such as <u>1K2N</u> (1.5%) when the nucleus had divided before the kinetoplast and cells with other abnormal K/N ratios including cells with more than 2K and/or 2N annotated <u>XKXN</u> (1.6%). As previously mentioned, when INb48::3cMyc is expressed in trypanosome cells, they exhibit detached flagella (18.3%). After 24hpi (+Tet 24h), this profile continued to augment with a further reduction in 1K1N (44.20%) and a further increase in 2K2N cells (27.5%). Additionally, the number of <u>1K2N</u> (abnormal) cells increased to 4.3% and 5.9% of the population showed XKXN phenotype. Finally, the number of cells with detached a flagella had also increased to 35.4%. INb48::3cMyc expressing cells in the 2K2N stage were further investigated and categorized according to linear organization of K and N from posterior to anterior orientation (Figure 7B). For non-induced cells (-Tet), most 2K2N trypanosomes had the normal KNKN organization with both flagella attached along the cell body (green bar). At 12hpi, the number of normal 2K2N population has decreased (from 93.5 to 45.8%) with the appearance of RNAi *Tb*BILBO1 knockdown-like 2K2N cells. 36.44% of the +Tet 12h total 2K2N population had detached flagella and an extended posterior end, which also included 15.8% with a K_NKN organization (here the underscore refers to an extended end, - orange bar). 13.1% of the abnormal population had a K_KNN kinetoplast organization (red bar) and 7.6% had a poorly segregated nuclei K_NKN* (yellow bar). The other 2K2N population 17.7% (blue bar) included cells that appeared normal in kinetoplast and nuclear organization KNKN but with detached flagella, and cells with oddly misplaced kinetoplast (KKNN) with detached flagella. At 24hpi (+Tet 24h), the proportion of 2K2N “normal-like” population was significantly reduced to 20.3% at the expense of abnormal cells mentioned above. The rest of the abnormal cells possessed a detached new flagellum which were distributed over a number of arrangements of K and N plus an extended posterior end K_KNN (red bar), K_NKN (orange bar) and K_NKN* (yellow bar) or without extended posterior ends (blue bar). These abnormal phenotypes are a clear indication of disruption of the cell cycle. As with RNAi knockdown of *Tb*BILBO1, it appears that there is a general cell cycle arrest at the 2K2N stage, indicating the cells cannot undergo division. Overall, this indicates that the DNA containing structures of the cell were able to divide and indeed separate (albeit imperfectly in some cases), but the trypanosome cell itself was unable to undergo cytokinesis. With regards to these cell cycle counts, similar findings were observed with RNAi knockdown of *TbBILBO1* (14), suggesting that the crucial steps and structures required for cytokinesis, were impaired after INb48::3cMyc expression. Taken together, this data shows that INb48 targeting of *Tb*BILBO1 can be as effective as RNAi knockdown.

**Fig 7.**
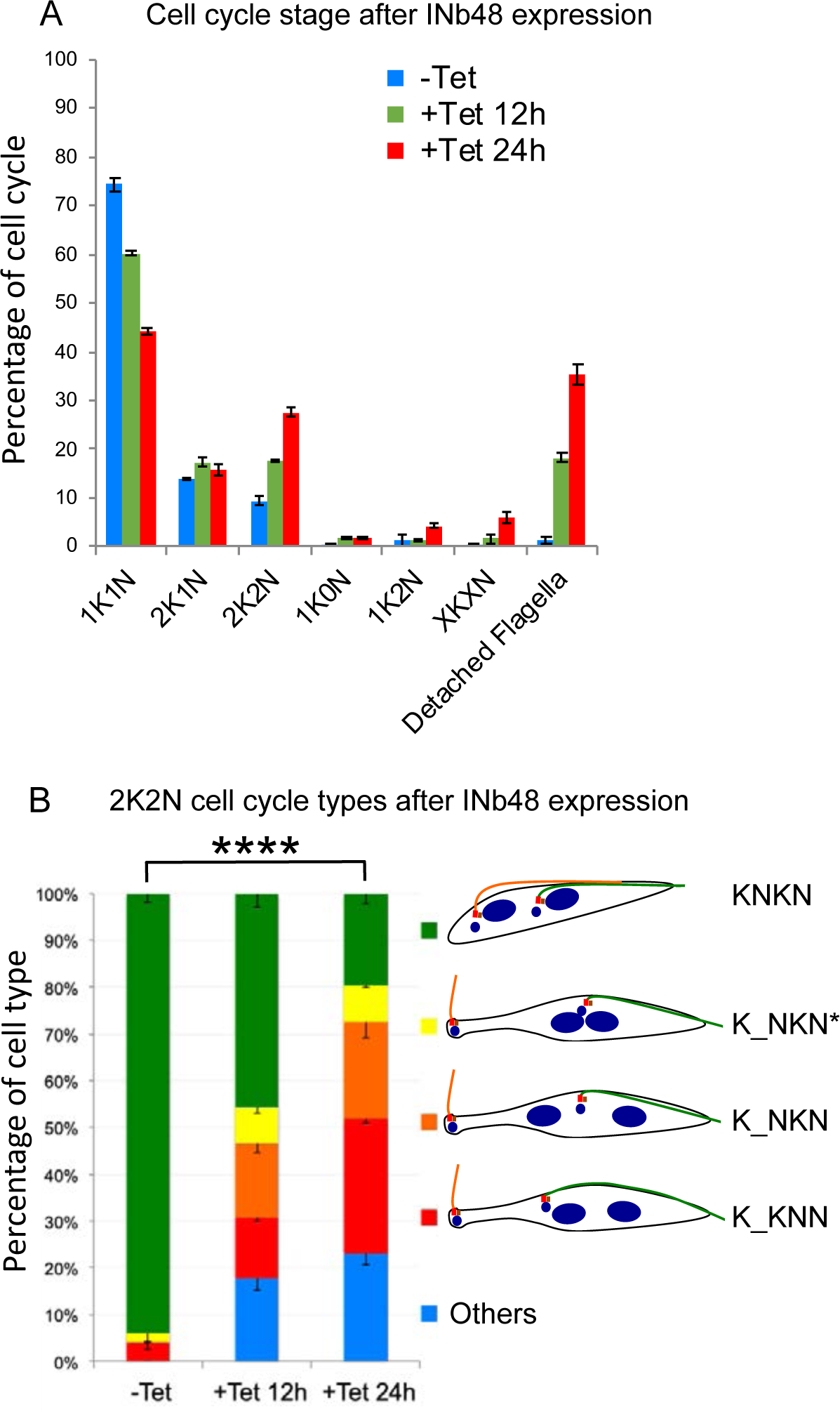
Expression of INb48::3cMyc disrupts the cell cycle in *T. brucei*. (A) INb48::3cMyc expression disrupts cytokinesis in *T. brucei.* Bar graph to show the percentage of trypanosomes counted at each stage of the cell cycle for non-induced (-Tet, blue bars), 12 hpi (+Tet 12h, orange bars) and 24 hpi (+Tet 24h, red bars) for INb48::3cMyc expression. Note that at 24hpi approximately 35% of cells have detached flagella. INb48::3cMyc expression leads to a decrease in 1K1N and an increase in 2K1N, 2K2N, abnormal 1K2N and XKXN phenotypes. N=300 trypanosomes per time point and each time point has been completed in triplicate. (B) Distribution of 2K2N cells after INb48::3cMyc expression in *T. brucei* showing strong similarities to *Tb*BILBO1 RNAi knockdown. The 2K2N cells were subdivided into 5 categories according to the visual appearance of the cell *i.e.,* possessing a detached flagellum, an extended posterior end, abnormal positioning of the kinetoplast and nucleus. **** *p* < 0.0001.

### Nb48 binds to full-length *Tb*BILBO1, an EF-hand mutated version and the coiled-coil domain truncation, in a heterologous system

Expression of full-length and truncated forms of *Tb*BILBO1 in a U-2 OS heterologous system has been published previously (23). In summary, that work had demonstrated that expression of full-length *Tb*BILBO1 in mammalian cells induces the formation of linear polymers with comma and globular shaped termini. To determine if Nb48 can bind to *Tb*BILBO1 as an antibody probe in this heterologous system, we expressed full-length *Tb*BILBO1 in U-2 OS cells and probed detergent-extracted cells with the purified Nb48 (Nb48_::HA::6His_) and with anti-*Tb*BILBO1 (1–110) (Figure 8). Non-transfected, extracted U-2 OS cells were probed with anti-TbBILBO1 (1–110) and Nb48_::HA::6His_, and showed a negative signal (Figure 8A). When full-length *Tb*BILBO1 was expressed it induced the formation of long BILBO1 polymers that were co-labelled with Nb48_::HA::6His_ and anti-*Tb*BILBO1 (1–110) (Figure 8B). *Tb*BILBO1 possesses two canonical calcium-binding EF-hand domains; 1 and 2. Mutation of EF-hand domain 1 alone, or both EF-hands together prevented the formation of linear polymers, however mutation of EF-hand domain 2 alone (Mut2 EF-hand) resulted in the formation of helical polymers (23). We therefore used Mut2 EF-hand to test for co-labelling with Nb48. Excellent co-labelling was observed when the Mut2 EF-hand was expressed: the helical polymers of the mutated BILBO1 co-label with Nb48_::HA::6His_, (Figure 8C), suggesting that mutating the calcium-binding site, and preventing calcium binding, does not affect Nb48 binding. We next expressed the *Tb*BILBO1 truncations T1, T2, T3, T4 (refer to Figure 2F) in U-2 OS cells and used the previously anti-*Tb*BILBO1 monoclonal IgM antibody to label T3 and T4 constructs (23). Previous studies had shown that *Tb*BILBO1 truncations T1 (aa1-170) and T2 (aa1-250) are soluble in U-2 OS cells and do not form polymers (23). No signal was observed when these extracted cells were probed by anti-*Tb*BILBO1 (1–110), furthermore no Nb48_::HA::6His_ signal was observed, (Figures 8D and E). Truncations T3 (aa171-587) and T4 (aa251-587), however, form long polymers when expressed in U2-OS cells and, as with FL polymers, these were positively co-labelled when probed with *Tb*BILBO1 and Nb48_::HA::6His_ (Figures 8F and G). As such, this suggest that the Nb48 epitope lies between the amino acids 251-587. These results illustrate that as a specific nanobody probe, Nb48_::HA::6His_, in a heterologous environment, performed equally to a rabbit polyclonal antibody (anti-*Tb*BILBO1 1-110) and to anti-*Tb*BILBO1 IgM.

**Fig 8.**
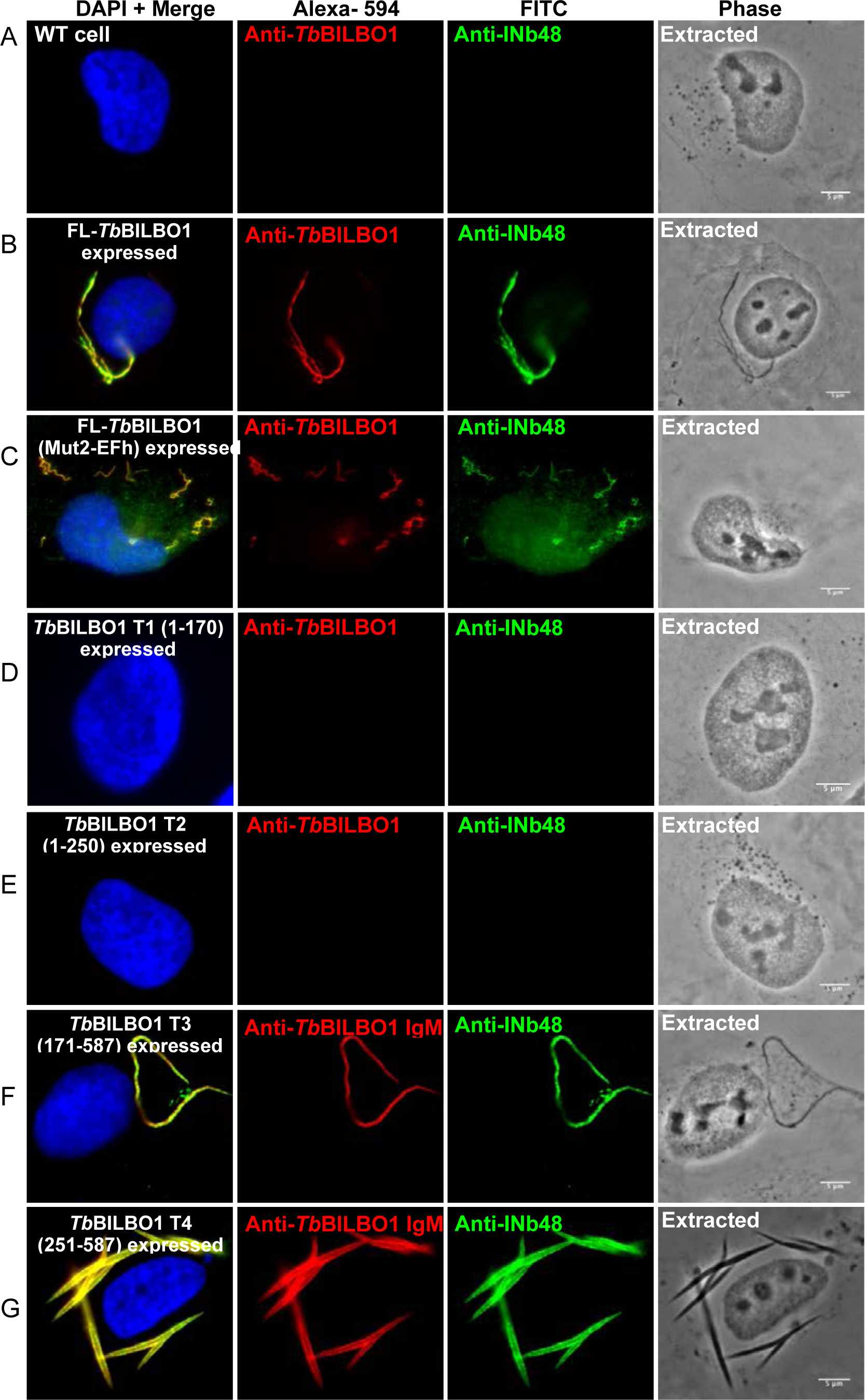
Nb48 recognizes and binds to *Tb*BILBO1 when expressed in a heterologous system. (A) Illustrates a non-transfected U-2 OS cell extracted with Triton X-100, probed with anti-*Tb*BILBO1 (1-110 rabbit) and purified Nb48_::HA::6His_ and revealed independently by anti-HA (mouse) and their respective secondary fluorophores: anti-rabbit, Alexa fluor 594 and anti-mouse, FITC-conjugated. No labelling is observed. (B) Illustrates a U-2 OS cell that is expressing FL *Tb*BILBO1 and probed as above. *Tb*BILBO1 forms polymers and Nb48_::HA::6His_ co-localises with these polymers. (C) Illustrates a U-2 OS cell expressing mutated EF-hand domain 2 (Mut2EF-*Tb*BILBO1) and probed as above. The typical spiral polymers of this mutant are formed and are co-labelled with rabbit 1-110 and Nb48_::HA::6His_. (D and E) Illustrate *Tb*BILBO1 truncations T1 (aa 1-170) and T2 (aa 1-250) which are both cytoplasmic after detergent extraction and are negative for anti-*Tb*BILBO1 and Nb48_::HA::6His_. (F and G) Illustrate the typical signals observed when *Tb*BILBO1 truncations T3 (171-587 aa) and T4 (251-587 aa) are expressed in U-2 OS; the resulting polymers formed and are both positive for *Tb*BILBO1 (in this case anti-*Tb*BILBO1 mouse monoclonal, IgM) and Nb48_::HA::6His_ (anti-HA rabbit, IgG). The IgM was used here because it binds to the C-terminus whereas anti-*Tb*BIL1BO (1–110) was raised to, and only binds to, the N-terminus of *Tb*BILBO1. Scale bar = 5μm. For the sake of convenience, the protein being probed is named as anti-Nb, plus the relevant nanobody name rather than its tag.

### Binding of INb48 to *Tb*BILBO1 in a heterologous system alters polymer formation

To determine if INb48 can bind to *Tb*BILBO1 in the U-2 OS heterologous system and influence polymer formation, we co-expressed INb48::3HA with full-length *Tb*BILBO1, (FL), Mut2 EF-hand, or truncated forms of *Tb*BILBO1 (Figure 9). Expression of INb48::3HA alone produces a cytoplasmic signal in whole cell preparations (Figure 9A), that was eliminated after detergent extraction (Figure 9B), demonstrating that INb48::3HA does not bind to any cytoskeletal structure in U-2 OS cells, and does not form polymers. When INb48::3HA was co-expressed with FL-*Tb*BILBO1, and cells detergent-extracted, a strong co-localisation was observed between the *Tb*BILBO1 signal and that of INb48::3HA, demonstrating that INb48::3HA was expressed, and binds to *Tb*BILBO1, *in vivo,* in detergent extracted cells (Figure 9C and D). Importantly, however, the typical polymer structures that *Tb*BILBO1 normally forms in U-2 OS cells (refer to Figure 8B) were absent. Instead, dense compacted structures, attached to thin minor polymers were formed. Similar dense structures were observed when Mut2-EFh was co-expressed with INb48::3HA, indicating that the EF hand domain 2 mutation (which prevents calcium binding) did not influence binding of INb48::3HA to *Tb*BILBO1 (Figure 9E). This data indicate that binding of INb48::3HA to *Tb*BILBO1 modifies and reduces linear polymerization in this environment.

**Fig 9.**
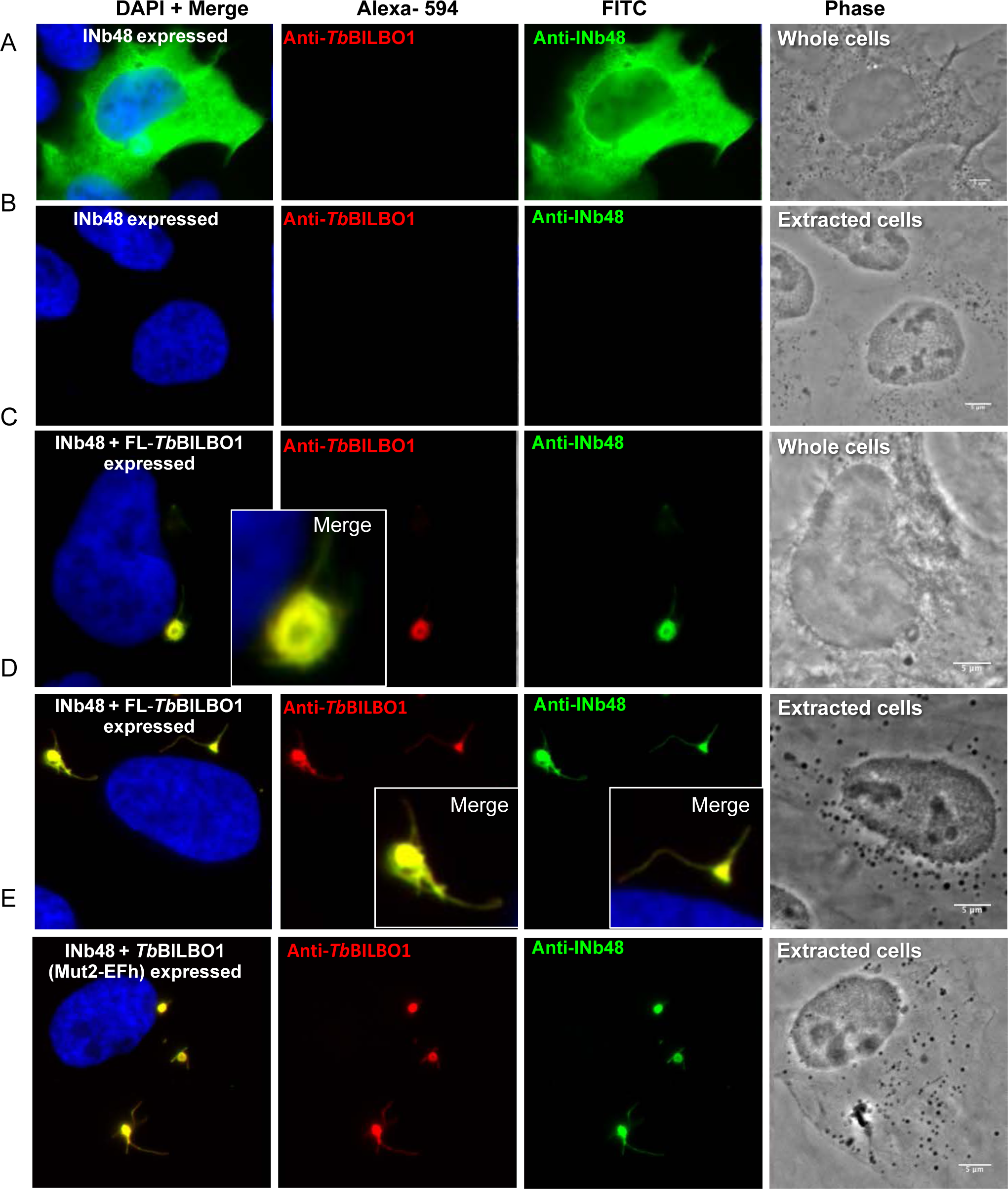
INb48::3HA interferes with *Tb*BILBO1linear polymer formation when co-expressed in U-2 OS cells In all experiments INb48, *Tb*BILBO1, *Tb*BILBO1 mutations and truncations were expressed in U-2 OS cells for 24h then processed for IFA. (A) IFA of U-2 OS cell expressing INb48::3HA only, and probed with anti-*Tb*BILBO1 and anti-HA to detect the intrabody. This cell shows that INb48::3HA is cytoplasmic. (B) Illustration of a cell expressing INb48::3HA only and probed with anti-*Tb*BILBO1 and anti-HA after detergent extraction, showing that INb48::3HA has been removed after extraction. (C) Illustration of a cell expressing INb48::3HA and FL-*Tb*BILBO1 simultaneously, and probed as in (A and B). It illustrates that INb48::3HA binds to the *Tb*BILBO1 polymers. However these polymers are different to WT polymers shown in Figure 8B, suggesting that INb48::3HA affects normal *Tb*BILBO1 polymer formation. (D) Illustrates a cell treated as in (C) but detergent extracted, demonstrating that INb48::3HA is not removed upon detergent extraction. (E) The cell in this image is simultaneously expressing an EF-hand domain 2 mutated form of *Tb*BILBO1 (Mut2EFh) and INb48::3HA. The typical helical spirals formed by the Mut2-EFhand protein are not formed when bound by INb48::3HA, suggesting again that INb48::3HA binding affects polymer formation. For the sake of convenience, the protein being probed is named rather than the tag. Scale bar = 5μm.

When INb48::3HA was co-expressed with T1 and with T2 (both have been demonstrated to be cytoplasmic - (23), a cytoplasmic signal was observed (Figures 10A and B). In contrast to the long, linear, polymers observed when T3 alone is expressed (Figure 8F), co-expression of T3 and INb48::3HA induced dense globular structures (Figure 10C) similar to the observations made when INb48::3HA was co-expressed with FL-*Tb*BILBO1 (Figures 9D) or Mut2-EFh (Figure 9E). Co-expression of INb48::3HA with T4 resulted in long polymers similar, but not identical, to expression of T4 alone (as observed in Figure 8G). However, the INb48::3HA bound T4 polymers were associated with many small, detergent insoluble, network-forming, polymers (Figure 10D), which were never observed in T4 only expression as previously described (14) (Figure 8G) and suggests that binding of INb48::3HA to T4 modifies normal polymerization. Taken together these data suggest a critical disturbance by INb48 in *Tb*BILBO1 polymer organization when co-expressed in a heterologous mammalian system. IFA controls for INb48::3HA and *Tb*BILBO1 expression in U-2 OS cells alone and together can be found in Supplementary data Figures 3A-D demonstrating a lack of labeling when no primary antibodies were used. Supplementary data Figure 4A-E shows the control intra-nanobody against GFP (INbGFP) expressed in U-2 OS cells exhibiting a cytoplasmic signal which was extracted in cytoskeleton preparations and did not co-localize or bind to *Tb*BILBO1 polymers or any other structure; negative and no primary antibody controls are also shown.

**Fig 10.**
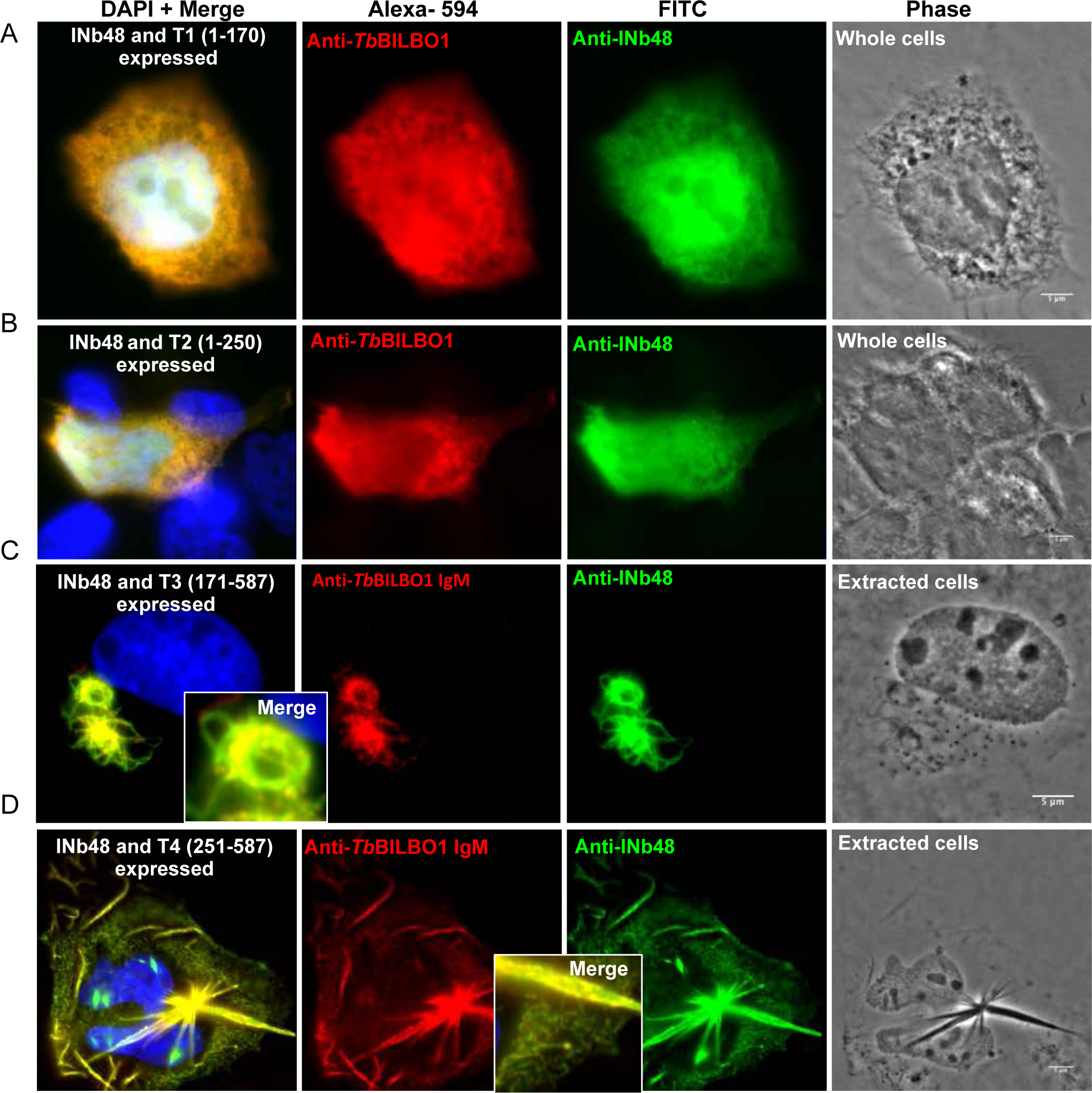
INb48::3HA expression in U-2 OS cells modifies polymers formed by *Tb*BILBO1 truncations, T3 and T4. As with the experiments of Figure 9, *Tb*BILBO1 proteins and INb48::3HA were expressed in U-2 OS cells for 24h; in this case INb48::3HA, was co-expressed T1-T4 truncations. Cells were then processed for IFA using either the anti-*Tb*BILBO1 (1–110) rabbit polyclonal, raised to and binding only, the N-terminus of *Tb*BILBO1, for detecting T1 and T2, or an in-house made anti-*Tb*BILBO1 IgM mouse monoclonal, which recognizes the C’ terminus of *Tb*BILBO1; anti-HA was used to detect INb48::3HA followed by their respective secondary antibodies. (A) Illustrates an IFA image of a U-2 OS cell simultaneously expressing INb48::3HA and *Tb*BILBO1 T1. These images illustrate that INb48::3HA and T1 are both cytoplasmic and therefore give an overlapping signal. (B) Illustrates an IFA image of a U-2 OS cell simultaneously expressing INb48::3HA and *Tb*BILBO1 T2. Similar to (A) both of these proteins are cytoplasmic and give an overlapping signal. (C) This is a IFA image of a U-2 OS cell simultaneously expressing INb48::3HA and *Tb*BILBO1 T3. The typical linear filaments formed by T3 are modified when bound by INb48::3HA, suggesting the intrabody binding inhibits the typical polymer formation. (D) Illustrates detergent extracted IFA images of U-2 OS cells simultaneously expressing INb48::3HA and *Tb*BILBO1 T4. The typical spindle shaped polymers, as observed in Figure 8G, are observed to a lesser extent but numerous smaller polymers are present in the cytoplasm and these are never present in cells expressing T4 alone. This suggests that the binding of INb48 to T4 modifies the polymer forming capacity of this truncation. Scale bar = 5μm. For the sake of convenience, the protein being probed is named as anti-Nb, plus the relevant nanobody name rather than its tag.

## Discussion

Nanobodies are rapidly being developed, not only as revolutionary therapeutics but also as biological tools. In mammalian cells they have been used to assess protein function intracellularly (intrabodies), explore protein-protein interactions and as protein inhibitors (42), (76). Nanobodies have been used in a number of experiments associated with trypanosomes including the potential of nanobodies as diagnostic tools, in experimental therapy, and as trypanolytic agents (40, 53, 60, 62, 64, 66–69, 77, 78). However, intrabodies have not been used to target the trypanosome cytoskeleton intracellularly as a knockdown tool. This study shows that Nb48 proved to be an excellent tool for detecting its trypanosomatid target antigen *Tb*BILBO1 by immunofluorescence (IFA) on fixed trypanosomes and by immuno-electron microscopy. Further, when expressed as an intrabody INb48 targeted specifically *Tb*BILBO1 and functioned to knockdown this essential cytoskeletal protein.

### INb48 modifies TbBILBO1 polymers in U-2 OS cells and in T. brucei

Nb48 and INb48 both bind to the T4 domain of *Tb*BILBO1 (aa 251-587) demonstrating that they recognize the same epitope *in vitro* and *in vivo*. Binding of INb48 to *Tb*BILBO1 in U-2 OS cells modifies polymer structure and induced condensed structures (Figure 9C -E) compared to the long, comma or annular shaped polymers that are normally observed (23). INb48 binds to the domain of *Tb*BILBO1 that includes the CCD and the LZ domains both involved in protein-protein interaction, and importantly, the CCD domain is required for dimerization and the LZ for polymerization (22, 23). Based on the structures observed in the U-2 OS cells, both dimerization and polymerization occur, but it is difficult to speculate on how INb48 affects the polymers. However, one could expect the same effect in parasites, resulting in alteration in FPC formation. Indeed, the expression of INb48 in *T. brucei* led to perturbation of the structure of the FPC and the HC, as seen by IFA and iEM labelling of *Tb*BILBO1 and *Tb*MORN1 respectively. INb48 labelled strongly the MtQ and the typical annular FPC structure and “hook and shank” aspect of the HC was not observed in many salt-extracted flagella in iEM experiments. Also, IFA experiments demonstrated (*via* the elongated structures labelled with an anti-BILBO1 polyclonal), that INb48 affects the biogenesis of the FPC. This suggests that *Tb*BILBO1 and *Tb*MORN1 traffic along the MtQ. The simplest explanation would be that INb48 blocks either the dimerization or the polymerization of *Tb*BILBO1 and this influences the formation of the FPC leaving *Tb*BILBO1 bound to the MtQ.

### INb48 induces BILBO1-RNAi knockdown-like phenotypes

The perturbations of *Tb*BILBO1 assembly by INb48 leads to RNAi knock-down phenotypes. The accumulation of intracellular vesicles following INb48 expression (Figure 6) indicates a disruption in endo-/exocytosis and is a consequence of the disturbance of the flagellar pocket and the HC, which was observed when *Tb*BILBO1 was knocked-down (14). However, INb48 does not have nuclear or ER targeting / retention signals which are sometimes used on INbs (79) supporting the idea that INb48 perturbs directly the assembly of *Tb*BILBO1 and the formation of the FPC. This would intimate that INb48 has its knockdown effect by either (a) binding directly to *Tb*BILBO1 during translation (where an IFA signal might be too weak to visualize), (b) during trafficking of *Tb*BILBO1 to the FPC on the MtQ and preventing formation of the FPC, (c) binding of INb48 on the older FPC and disrupting duplication (if we consider a semi-conservative hypothesis for FPC formation), (d) a combination of these events. INb48 was observed at the FPC as early as 3hrs of induction (Sup. Figure 1B; 3h induction) making it difficult to determine exactly where in the cell the knockdown effect occurs. An identical version of INb48 (with a single amino acid mutation of CDR3 was observed on the FPC as early as 15min of induction – not shown). Considering hypotheses (a) and (b) from above, it could be speculated that binding to *Tb*BILBO1 as it is trafficked on the MtQ in the cytoplasm or on the ribosome might prevent chaperone binding, or modify *Tb*BILBO1 three-dimensional structure resulting in degradation. We are currently investigating the knockdown effect in more detail but since we hypothesize that the MtQ are used for trafficking of FPC proteins, then transport of INb48 to the FPC would in this case most likely occur through binding to *Tb*BILBO1 in the cytoplasm and or transfer onto the MtQ. This may also answer, in part, the question of how the FPC, a flagellum-associated complex of proteins, is built.

### Conclusion

RNA interference (RNAi) has been used to greatly enhance knowledge of gene and consequently protein function in living cells. However, RNAi can have several limitations, such as incomplete knockdown of the target protein. Furthermore, the double-stranded RNA produced during RNAi has a relatively short half-life compared to an intrabody. One advantage of intrabodies is that they act at a post-translation stage, therefore they can target a specific isoform or post-translational modification of a protein (78). Also, in case of essential or structural proteins, induction of the expression of INbs at a specific cell-cycle, life-cycle stage or point in organelle biogenesis may provide very subtle and precise phenotypes. In this regard, nanobodies also have advantages over conventional antibodies due to their smaller size, allowing access to cryptic epitopes, and ability to cross the blood-brain barrier (81, 82). Conventional antibodies have a further disadvantage of multi-epitope cross-reaction, a phenomenon exemplified by IgE in allergen cross-reaction, whereas, nanobodies are, theoretically, single epitope specific (37, 38, 83).

Our research data support the hypothesis that targeting minor, yet essential, cytoskeletal proteins is of considerable merit in the search to understand parasite biology. They also suggest that the use of intrabodies might be useful in organisms that do not have RNAi machinery or to characterize the cell biology of emerging pathogens, marine organisms, protists and new model organisms such as those described by Faktorová *et al* (84). The data presented here illustrates for the first time a functional anti-cytoskeletal intrabody in *T. brucei*, that is able to precisely target and bind to its target protein epitope and induce disruption of a cytoskeletal structure leading to rapid cell death.

## Materials and Methods

### Ethics statement

The alpaca immunization was carried out by Nanobody Service Facility, VIB Rijvisschestraat 120, 9052, Belgium and was approved by the Ethical Committee for Animal Experiments of the Vrije Universiteit Brussel (VUB), Brussels, Belgium.

### Alpaca immunization and Nb library construction

HIS-tagged, full-length *T. b. brucei* BILBO1 protein (6HIS::*Tb*BILBO1) was expressed in bacteria, purified in urea on Ni-NTA resin by affinity column purification, as described in (25), and sent to Nanobody Service Facility (NSF, VIB, Brussels). The nanobody library was constructed as described by NSF-VIB (Nanobody Service Facility, Belgium) and produced 2×10^8^ independent transformants that were subjected to two rounds of panning, performed on solid-phase coated full-length *Tb*BILBO1 antigen (10μg/well). Ninety-five colonies from round two were randomly selected and analyzed by ELISA for the presence of antigen-specific Nbs in their periplasmic extracts. Thirty-four colonies scored positive, representing seven different Nbs from three different groups based on their gene sequences: Nbs in each group have very similar sequence data. The Nb genes were cloned into a pMECS vector containing a secretion PelB leader signal sequence at the N-terminus, a hemagglutinin A (HA) tag and a hexa-histidine (His) tag at the C-terminus, for purification and immunodetection. These vectors were transformed into TG1 *E. coli* and stored at -80°C.

### Nanobody cloning and bacterial expression

Nbs were sub-cloned from pMECS to pHEN6c vector as follows: TG1 *E. coli* harboring the Nbs, were grown overnight on solid agar + ampicillin (100μg/mL). The Nb genes were amplified by PCR directly on colonies using specific primers A6E (5’GAT GTG CAG CTG CAG GAG TCT GGR GGA GG 3’) and PMCF (5’ CTA GTG CGG CCG CTG AGG AGA CGG TGA CCT GGG T 3’). PCR fragments were digested with *Pst*I and *Bst*II then ligated into the pHEN6c vector harboured in XL1 blue *E. coli* and stored at -80°C. For expression and purification, WK6 *E. coli* were transformed with pHEN6c harbouring the Nb. PCR on transformed colonies, using M13R (5’ TCA CAC AGG AAA CAG CTA TGA C 3’) and PMCF (5’ CTA GTG CGG CCG CTG AGG AGA CGG TGA CCT GGG T 3’) confirmed the Nb insert. *Tb*BILBO1, *Tb*BILBO1 EF-hand domain mutant 2 and *Tb*BILBO1 truncations T1, T2, T3 and T4 were expressed in the heterologous mammalian system, U2-OS, by using pCDNA3 vector as described in (23). Nb48 sequence was cloned into the pcDNA3.1 C-terminus tag 3HA vector (derived from pCDNA3.1 Invitrogen) between *Nhe*I-*Xho*I restriction sites and expressed transiently as an intrabody, INb48::3HA.

### Intra-nanobody expression in Trypanosoma

The Nb inserts were digested out from pHEN6c with *Hind*III and *Nde*I and cloned into *Hind*III and *Nde*I digested pLew100-3cMyc (86). Plasmid sequences were confirmed by DNA sequencing.

### Nanobody expression and purification

Freshly transformed WK6 with pHEN6c-Nb vectors were inoculated into 10mL LB + ampicillin (100μg/mL) + 1% glucose and incubated overnight at 37°C, shaking at 250 rpm. Terrific broth (TB) medium was prepared for Nb expression (2.3g/L KH2 PO4, 16.4g/L K2 HPO4.3H2O, 12g/L Tryptone, 24g/L Yeast extract, 4% glycerol). One mL of culture was added to 330mL TB + 100μg/mL ampicillin, 2mM MgCl2 and 0.1% glucose, incubated at 37°C, shaking 250 rpm, until a mid-log phase of growth OD600 of 0.6-0.9 was reached. Nb expression was induced with 1mM Isopropyl β-D-1-thiogalacto-pyranoside (IPTG). The culture was incubated for 17 hours at 28°C with shaking. Nb was extracted from the periplasm as follows. Cultures were centrifuged for 16 minutes at 4,500 x g. The pellet was re-suspended in 4mL of TES (0.2M Tris pH8.0, 0.5mM EDTA, 0.5M sucrose), then shaken for 1 hour on ice at 110 rpm. Four mL of TES/4 (TES diluted 1:4 in water) was added and further incubated on ice for 1 hour with shaking, then centrifuged for 60 minutes at 4,500 x g at 4°C to releases the contents of the periplasmic space (87). The supernatant contains the periplasmic extract with Nb. A control periplasmic extraction was performed using the same procedure as above, without the Nb insert in the vector.

The periplasmic extract was filtered on 0.45 µm, and loaded onto a HisTrap^TM^FF column (GE Healthcare) pre-equilibrated with running buffer (PBS, 20mM imidazole, 10% glycerol, protease Inhibitor Cocktail Set III Calbiochem, Ref. 539134-1 at 1:10,000 dilution). The column was washed with five column volumes (CV) of lysis running buffer and bound protein was eluted by 500 m*M* imidazole/running buffer with 10 CV. Elutions containing the highest concentration of purified nanobody were pooled and dialysed against 1xPBS + 20% glycerol using Slide-A-Lyzer^TM^ (Thermo Scientific B2162132) and stored in 1xPBS + 20% glycerol. To verify the presence of Nb in purified elutions, 10μL from each elution was run on 15% SDS-PAGE gel stained with Coomassie Brilliant Blue (Instant Blue^TM^ Expedeon Ltd). Proteins from a second identically loaded gel were transferred to PVDF membrane using a semi-dry transfer method (BIORAD) and a western blot was performed using anti-HIS (Sigma H-1029, mouse, 1:3000) and anti-HA (Biolegend MMS-101R or, Santa-Cruz 7392, 1:1000) antibodies to detect the Nb, followed by anti-mouse HRP conjugated (Jackson, 115-035-044, 1:10,000 or, Jackson, 515-035-062 1:1000).

### Surface Plasmon Resonance Assays

Surface plasmon resonance (SPR) was used to measure the binding affinity of the nanobodies to *Tb*BILBO1 purified protein, the benefits being it is measured in real-time and is label-free. All solutions and buffers used were filtered and degassed. The experiments were performed at 25°C using a Biacore^TM^ T200 instrument (GE Healthcare Life Sciences, Uppsala, Sweden) flowed with EP+ buffer (10mM HEPES, pH 7.4, 150mM NaCl, 3mM EDTA, 0.05% Tween-20) from the manufacturer as running buffer. For *Tb*BILBO1 expression, 6His::*Tb*BILBO1 protein was overexpressed in bacteria, soluble supernatant purified on Ni-NTA resin as describe in (13). Briefly, the cells were grown at 37°C in Luria–Bertani (LB) medium containing 50µ g/mL kanamycin to OD600 0.6. The cells were cold-shocked on ice for 30 minutes and induced with of 1mM of IPTG for 2 hours at 16°C. After centrifugation, the cell pellet was lysed by sonication in lysis buffer (2mM Tris–HCl pH 7.4, 0.5M NaCl, 20mM imidazole, 10% glycerol, Protease Inhibitor Cocktail Set III Calbiochem, 1:10,000) and centrifuged (10,000g, 30 minutes, 4°C) to remove cell debris. The supernatant was filtered at 0.45µ m, and loaded onto a HisTrapTMFF column (GE Healthcare) pre-equilibrated with the same lysis buffer. The column was washed with 5 column volumes (CV) of lysis buffer and bound protein was eluted by 500mM imidazole with 10 CV in lysis buffer. *Tb*BILBO1 fractions were pooled and dialyzed against PBS, 10% glycerol and concentration was measured using Thermo Scientific™ NanoDrop 2000. *Tb*BILBO1, prepared at 50µ g/mL in 10mM sodium acetate buffer, pH 7, was immobilized on a one flow cell of a CM5 sensor chip (GE-Healthcare, lot no 10266084) by amine coupling as indicated by the manufacturer and the protein solution was injected for 11 minutes; 2973 resonance units (RU) of protein were immobilized. One flow cell was left blank for double-referencing of the sensorgrams. The nanobodies were dialyzed against the EP+ running buffer and were injected over the target by Single-Cycle Kinetics (SCK). This method consists of injecting the partner sequentially at increasing concentration without a regeneration step between each concentration injected. Protein concentrations were determined using a Thermofisher Nanodrop^TM^ One spectrophotometer (Ozyme) and were injected over the target by Single-Cycle Kinetics (SCK) (87, 88). The surface was regenerated after each SCK cycle with two 1 minute pulses of 10mM glycine-HCl pH 2.1. The sensorgrams were analyzed with Biacore T200-v2.0 evaluation software using a Langmuir 1:1 model of interaction to determine the association (k_a_) and dissociations (k_b_) rate constants. A regeneration test was performed using 10mM glycine-HCl pH 2.1 (GE healthcare), 10mM glycine HCl pH2.1, at 30μg/min for 30 seconds with *Tb*BILBO1 alone to ensure no loss of signal. The dissociation equilibrium constant, K_D_, was calculated as k_b_/k_a_. The values shown are the average and standard deviation of three independent experiments with samples injected in duplicate.

### Trypanosome cell lines, culture and transfection

Procyclic (PCF) cell line Tb427 29.13 *Trypanosoma brucei brucei* (86) was grown in SDM-79 medium (GE Healthcare, G3344-3005) pH7.4 supplemented with 2 mg/mL hemin (Sigma Aldrich, H-5533), 10% fetal bovine serum complement deactivated at 56°C for 30 minutes (FBS; Gibco, 11573397;) and incubated at 27°C. Selection antibiotics hygromycin (25μg/mL) and neomycin (10μg/mL) were added to the media to maintain the plasmid vectors pLew29 and pLew13 (86). For transfection, the pLew100_Nb_3cMyc plasmids were linearized with *Not*I, then concentrated and purified by ethanol precipitation. Laboratory cell lines PCF Tb427 29.13 were grown to 5-10x10^6^ cells/mL, then 3x10^6^ cells were electro-transfected with 10μg of plasmid using the program X-001 of the Nucleofector®II, AMAXA apparatus (Biosystems) as described in reference (86) with the transfection buffer described in (89). After transfection, clones were obtained by serial dilution and maintained in logarithmic phase growth at 2x10^6^ cells/mL . Selection antibiotics hygromycin (25μg/mL) and neomycin (10μg/mL) were added to the media to maintain the laboratory cell lines Tb427 29-13 (procyclic forms, PCF). Phleomycin (5μg/mL) was added to the media of transfected cells to select for those harboring the pLew100-3cMyc vector and the Nb gene. Expression of Nb in the parasites (intrabody) was induced with tetracycline at 0.1ng/mL - 1μg/mL tetracycline. Growth curves representing logarithmic number of cells were calculated by counting the number of cells every 24 hours using Mallassez counting chamber slides. Error bars represent SEM = standard error of the mean.

### Mammalian cell line culture and transfection

U-2 OS cells (human bone osteosarcoma epithelial cells, ATCC Number: HTB-96) (88) were grown in D-MEM Glutamax (Gibco) supplemented with final concentrations of 10% fetal calf serum (Invitrogen), 100 units/mL of Penicillin (Invitrogen), and 0.100mg/L of Streptomycin (Invitrogen) at 37°C plus 5% CO_2_. Exponentially growing U-2 OS cells in 24 well plate with glass coverslips were lipo-transfected as described in (89) with 0.5μg of DNA using lipofectamine 2000 in OPTIMEM (Invitrogen) according to the manufacturer’s instructions and processed for immunofluorescence 24 hours post-transfection.

### Western blot analysis

Trypanosomes were collected, centrifuged at 800 x g for 10 minutes, and washed in 1xPBS. For whole cell samples, the number of trypanosomes to be loaded per well were calculated, then re-suspended in protein sample buffer 2x, plus nuclease 250 IU/mL (Sigma Aldrich, Ref. E1014). The samples were boiled at 100°C for 5 minutes then stored at -20◦C until required. 15% SDS-PAGE gels were prepared and samples loaded at 5x10^6^ WT PCF trypanosome cells, or PCF expressing pLew100-Nb-3cMyc (induced with tetracycline 1μg/mL, 24 - 48 hours). For detection of Nb in bacterial periplasmic extract, 2x10^8^ bacterial cells or equivalent volume of periplasmic extract, flow through or elution were loaded. Samples were separated according to manufacturer’s recommendations then transferred in a semi-dry system (BIORAD Trans-Blot Semi-dry transfer cell 221BR54560) onto PVDF membrane and incubated with blocking solution (BS: 5% skimmed milk powder, 0.2% Tween 20 in 1xTBS) for 60 minutes. For detection of intrabody, INb::3cMyc, expression in trypanosomes, an anti-cMyc antibody (Sigma, rabbit, C-3956) was diluted in blocking solution 1:1000 and incubated with the membranes overnight at 4°C. Membranes were washed in 1xTBS and then in BS before probing with anti-rabbit antibody conjugated to Horse Radish Peroxidase (HRP) (1:10,000 in BS, Sigma, goat, A9169) for 60 minutes at RT. The following antibodies were also used as primaries: anti-*Tb*BILBO1 aa1-110 (rabbit polyclonal, 1:1,000) (23); anti-*Tb*Enolase (rabbit polyclonal, 1:25,000) (92), and anti-TAT1 (anti-Tubulin mouse monoclonal, 1:1,000) (93) were used as loading controls. A secondary anti-mouse HRP antibody was also used, diluted 1:10,000 (Jackson, sheep, 515-035-062). Revelation was carried out using Image Quant LAS 4000 (GE) or Chemidoc (Biorad). Purified Nbs were used as primary antibodies in western blot assays to detect the presence of *Tb*BILBO1 in whole trypanosome cells. Whole cell extract of 5x10^6^ WT PCF trypanosome cells, or PCF expressing *Tb*BILBO1-3cMyc, full-length or truncations of *Tb*BILBO1 (T1, T2, T3, T4, induced with tetracycline 1*μ*g/mL, 24 hours), were loaded on 12 or 15% SDS-PAGE. Western blots were carried out as described above and the purified Nb::3HA::6His was used at a concentration of 10ug/mL to probe the membranes. To detect purified nanobody, an anti-HA (1:1000, Biolegend, 901513) or anti-HIS (1:10,000, Sigma H1029) was used, followed by an anti-mouse HRP conjugated secondary (above).

### Immunofluorescence assay (IFA)

One mL of log-phase trypanosomes were collected, centrifuged for 5 minutes at 1000 x g, washed in 1x PBS, then 20μL was loaded onto poly-L-lysine 0.1% solution (P8920 Sigma-Aldrich) coated slides for 4 minutes to adhere. Whole cells were fixed in methanol at -20°C for at least 60 minutes. Cytoskeleton extraction was with 0.25% NP40 in PIPES buffer (100 mM PIPES pH 6.8, 1mM MgCl2) for 5 minutes, washed twice in PIPES buffer and cytoskeletons were fixed with 1% PFA or methanol. Fixed trypanosomes were incubated with single or combination antibodies: anti-cMyc (monoclonal antibody IgG1 9E10, 1:20) to detect expressed INb, anti-*Tb*BILBO1 (amino acids 1–110, rabbit, 1517028 bleed 1 1:4000), anti-*Tb*MORN1 (rabbit 1:5000), anti-*Tb*PFR2 (rabbit 1:200), anti-*Tb*BLD10 (rabbit 1:2000), for 60 minutes, washed twice in 1xPBS, then incubated with secondary antibodies: anti-mouse IgG FITC-conjugated (Jackson, 115-095-164), and anti-rabbit Alexa fluor 594-conjugated (Invitrogen, A-11012). Kinetoplasts and nuclei were stained with 10*μ*L DAPI (10*μ*g/mL) in 1xPBS for 2 minutes and slides were mounted with Slowfade® Gold antifade reagent (Invitrogen, Molecular Probes). Images were acquired on a Zeiss Axioimager Z1 using MetaMorph software and a Roper CoolSNAP HQ2 camera. For Nbs used as probes in IFA of fixed trypanosomes, PCF Tb427 29.13 wild type (WT) and *Tb*427 29.13 pLew100-*Tb*BILBO1-3cMyc were fixed (as above) incubated with Nb (0.25mg/mL) for 1 hour, followed by anti-HA (Biolegend, 901513, mouse) diluted 1:200. For co-labelling with *Tb*BILBO1, anti-*Tb*BILBO1 (1–110) was used diluted 1:4000 in 1xPBS. Secondary antibodies were used with 1:100 dilution: anti-mouse FITC-conjugated (Sigma, F2012), anti-rabbit Alexa fluor 594-conjugated (Invitrogen, A11012). For IFA on U-2 OS cells, cells were fixed and processed as described in (23, 25). The primary antibodies anti-*Tb*BILBO1 (1–110), 1:4000 dilution; anti-*Tb*BILBO1 (IgM) mouse monoclonal, 5F2B3, (14) 1:2 dilution; anti-HA epitope tag (mouse monoclonal, Biolegend, 1:1000); anti-HA (rabbit, GeneTex GTX115044, 1:1000) were incubated for 1h in a dark moist chamber. After two 1 x PBS washes, cells were incubated for 1 hour respectively with the secondary antibodies goat anti-rabbit IgG (H+L) conjugated to Alexa fluor 594 (Molecular Probes A-11012, 1:400); goat anti-mouse IgM conjugated to Alexa fluor 594 (Invitrogen, 1:400); goat anti-mouse whole molecule conjugated to FITC (Sigma F-2012, 1:400); goat anti-rabbit IgG (H+L) conjugated to Alexa fluor 594 (Molecular Probes A-11012, 1:400). Purified Nb48_::HA::6His_ raised against *Tb*BILBO1 was used as a probe (10 μg/mL) and incubated for 1h in a dark moist chamber. After two PBS washes, cells were incubated for 1 hour with the secondary antibody directed against the HA tag, anti-HA epitope tag (Biolegend, 1:1000) then washed twice in PBS and incubated 1 hour with the tertiary antibody goat anti-mouse whole molecule conjugated to FITC (Sigma, 1:400). The intrabody anti-GFP nanobody 3 x HA-tagged (INbGFP::3HA) was probed with anti-HA mouse IgG Biolegend, diluted 1:1000 in PBS FCS 10%, saponin 0.1% and probed with anti-mouse goat whole molecule FITC, Sigma, diluted 1:100 in PBS FCS 10%, saponin 0.1%. The nuclei were stained with DAPI (0.25 μg/mL in PBS for 5 minutes). Slides were washed and mounted with Prolong (Molecular Probes S-36930). Images were acquired on a Zeiss Imager Z1 microscope with Zeiss 63x objective (NA 1.4), using a Photometrics Coolsnap HQ2 camera and Metamorph software (Molecular Devices), and processed with ImageJ (NIH).

### Flagella preparation for IFA

Ten mL of mid-log PCF cells were collected and washed once in 1xPBS for 5 minutes at 1000 x g at RT. Cells were extracted with 1mL 1% NP40 in 100mM PIPES pH6.9, 1mM MgCl2 for 7 minutes at RT to make cytoskeletons. Cytoskeletons were centrifuged for 10 minutes at 5,000 x g at 4◦C and further extracted for 15-30 minutes on ice with 1% NP40 in 100mM PIPES buffer + 2mM MgCl2, containing 1M KCl. Flagella were centrifuged for 5 minutes at 5,000 x g at 4◦C, the supernatant was removed and flagella were resuspended in 500μL PIPES buffer. Flagella were washed twice in 100mM PIPES buffer + 1mM MgCl2 for 5 minutes 5000 x g at 4◦C before being deposed on polylysine coated slides and left to adhere for 5–10 minutes. The flagella were fixed at -20◦C in MeOH or in 3% PFA as described above.

### Transmission Electron microscopy

Ten mL of mid-log phase procyclic, *Tb*427 29.13 wild type (WT) cells, pLew100-Nb48-3cMyc, (induced for 24h) were fixed, dehydrated and embedded, cut and stained as in (13) with the exception that tannic acid was omitted and cells were initially fixed in their culture medium by the addition of glutaraldehyde to a final concentration of 2.5% and paraformaldehyde at 3.7% for two hours then pelleted at 1,000 x g and fixed for 2 hours in 2.5% glutaraldehyde in phosphate buffer pH 7.4.

### Immuno-electron microscopy

Ten mL of mid-log phase ∼ 5 x 10^6^/mL pLew100-Nb48-3cMyc cells were induced with 1*μ*g/mL tetracycline for 24 hours. Five mL of induced cells were harvested at 1,000 x g, for 5 minutes, washed twice with PBS by centrifugation (1,000 x g) resuspended in 500µ L of PBS. Freshly glow-discharged, formvar and carbon-coated G2000-ni nickel grids (EMS) were floated on the droplets and the cells allowed to adhere for 15 minutes. Grids were then moved onto a drop of 1% NP-40 in PEME buffer (100 mM PIPES pH 6.8, 1 mM MgCl2, 0.1mM EGTA) (5 min, RT), followed by incubation on a 500µ L drop of 1% NP-40, 1M KCl in PEME buffer for 15 minutes (3 x 5 minutes, 4^0^C). Flagella were washed twice (2 x 5 minutes) in PEME buffer at RT, equilibrated and blocked on 50μL drops of 2% fish skin gelatin (Sigma-Aldrich G7041) or 0.5% BSA, 0.1%tween 20 in PBS, incubated in 25μL of primary antibody diluted in blocking buffer (each primary antibody was used either alone, or mixed with a second primary antibody), 1 hour at room temperature (RT). Anti-cMyc IgG mouse monoclonal antibody, (9E10, purified, a kind gift from Pr. Klaus Ersfeld), was diluted 1:10 or protein G purified/concentrated supernatant (diluted 1:50). Anti-*Tb*BILBO1 (1–110) was diluted 1:200 or 1:400. Grids were blocked and incubated in secondary antibody for 1 hour at RT (anti-mouse goat GMTA 5nm gold, and/or anti-rabbit goat 15nm gold GAR15, BBInternational) 1:10 in 0.2% fish skin gelatin in PBS. Grids were blocked, and fixed in 2.5% glutaraldehyde in 0.2% fish skin gelatin in PBS. Samples were negatively stained with 1% aurothioglucose (Sigma-Aldrich) (10μL for 30 seconds). Micrographs were taken on a Phillips Technai 12 transmission electron microscope at 100 kV.

## Acknowledgments

We thank our lab members for insightful comments on the manuscript, Dr. Brooke Morriswood, (University of Wurzburg, Germany) for the anti-*Tb*MORN1 antibody, Pr. Keith Gull (University of Oxford, England) for the anti-TAT1 (anti-Tubulin) antibody, Dr. Zyin Li (University of Texas, USA) for the anti-*Tb*BLD10 antibody, Pr. Klaus Ersfeld (University of Bayreuth, Germany) for the anti-cMyc (9E10) antibody, Dr. Frédéric Bringaud, (University of Bordeaux, Bordeaux, France) for the anti-Enolase antibody and Dr. Nicolas Biteau (University of Bordeaux, Bordeaux, France) for the anti-*Tb*PFR2 rabbit polyclonal antibody. We acknowledge NSF, VIB (Nanobody service facility), Brussels, for producing the nanobodies. This work was supported by internal grants (CNRS and University of Bordeaux) and support from the ANR, LABEX ParaFrap, ANR-11-LABX-0024 awarded to D.R.R.

## Supporting Information

**Sup Fig. 1.**
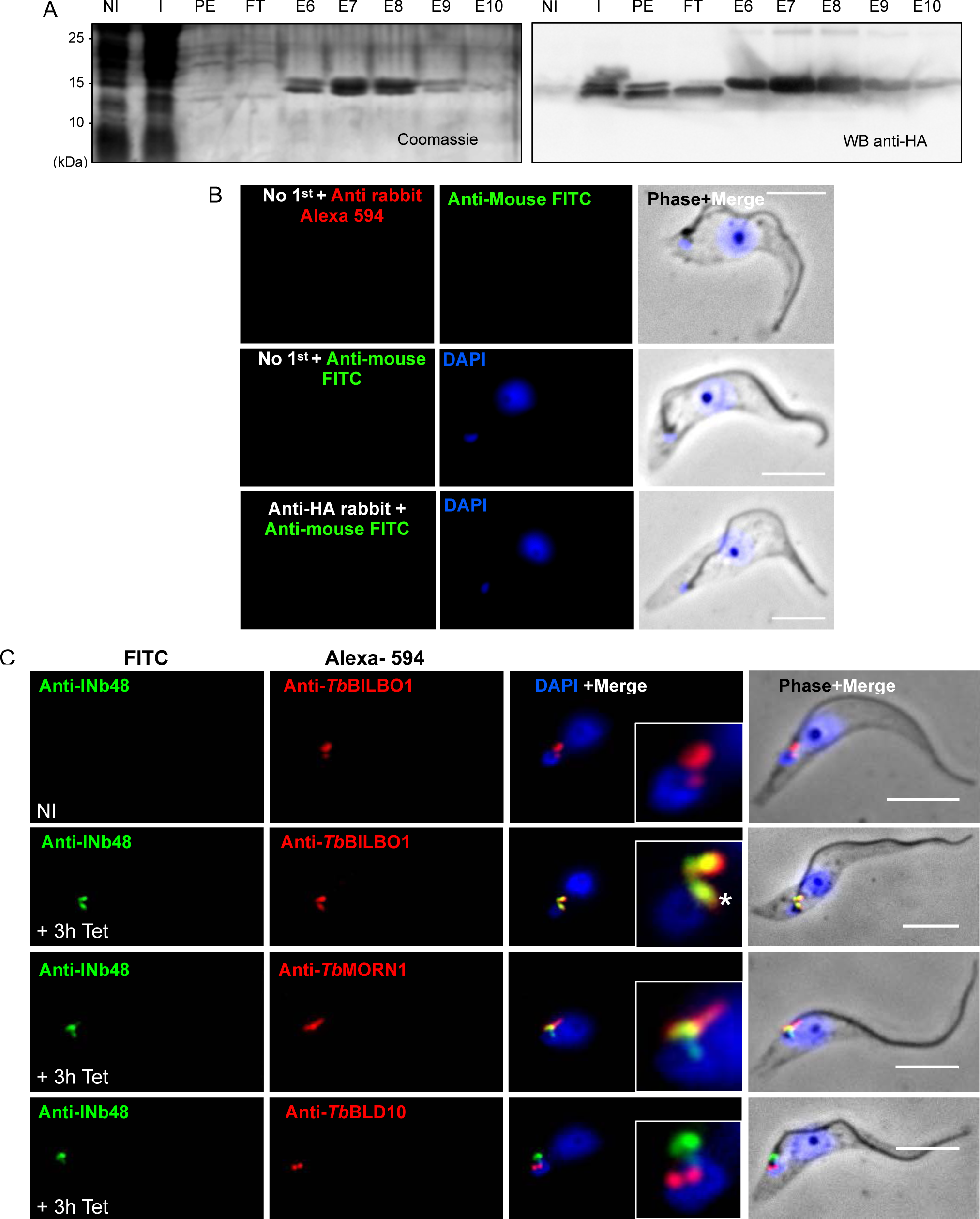
(A) Coomassie Blue 15% SDS-PAGE stained gel (left panel) and western blot (right panel) of the periplasmic extraction of Nb48_::HA::6His_ from WK6 *E. coli*. NI = non-induced bacteria; I = induced bacteria with 1mM IPTG; PE = periplasmic extract; FT = flow through; E = elutions from column. Load per well was 2x10^8^ bacterial cells or equivalent volume of PE/FT/E. (B) IFA controls for (Figure 2A-D and Figure 1D-G). First panel is a WT *T. brucei,* procyclic, cytoskeleton probed with anti-rabbit Alexa fluor 594 followed by anti-mouse FITC but no primary antibodies, demonstrating that there is no cross-reaction between these secondary antibodies or on the cytoskeleton. The second panel is a negative control of WT cytoskeleton probed with anti-mouse FITC but no primary antibody, which demonstrates that there is no cross-reaction between this secondary antibody and the cytoskeleton. The third panel is a negative control of a WT cytoskeleton probed with anti-HA rabbit followed by anti-mouse FITC, which demonstrates that there is no cross-reaction between these antibodies on the cytoskeleton. (C) In non-induced INb48::3cMyc cells (first panel), anti-cMyc reveals no labeling. After INb48::3cMyc expression is induced for 3 hours (+3h; second panel) labelling is visible at the FPC (green) co-localizing with *Tb*BILBO1 (red) at the FPC and along the MtQ (*). The two other panels show localization of INb48::3cMyc (green) with *Tb*MORN1 (red) and *Tb*BLD10 (a basal body marker; red) respectively. Scale bar = 5μm.

**Sup Fig 2.**
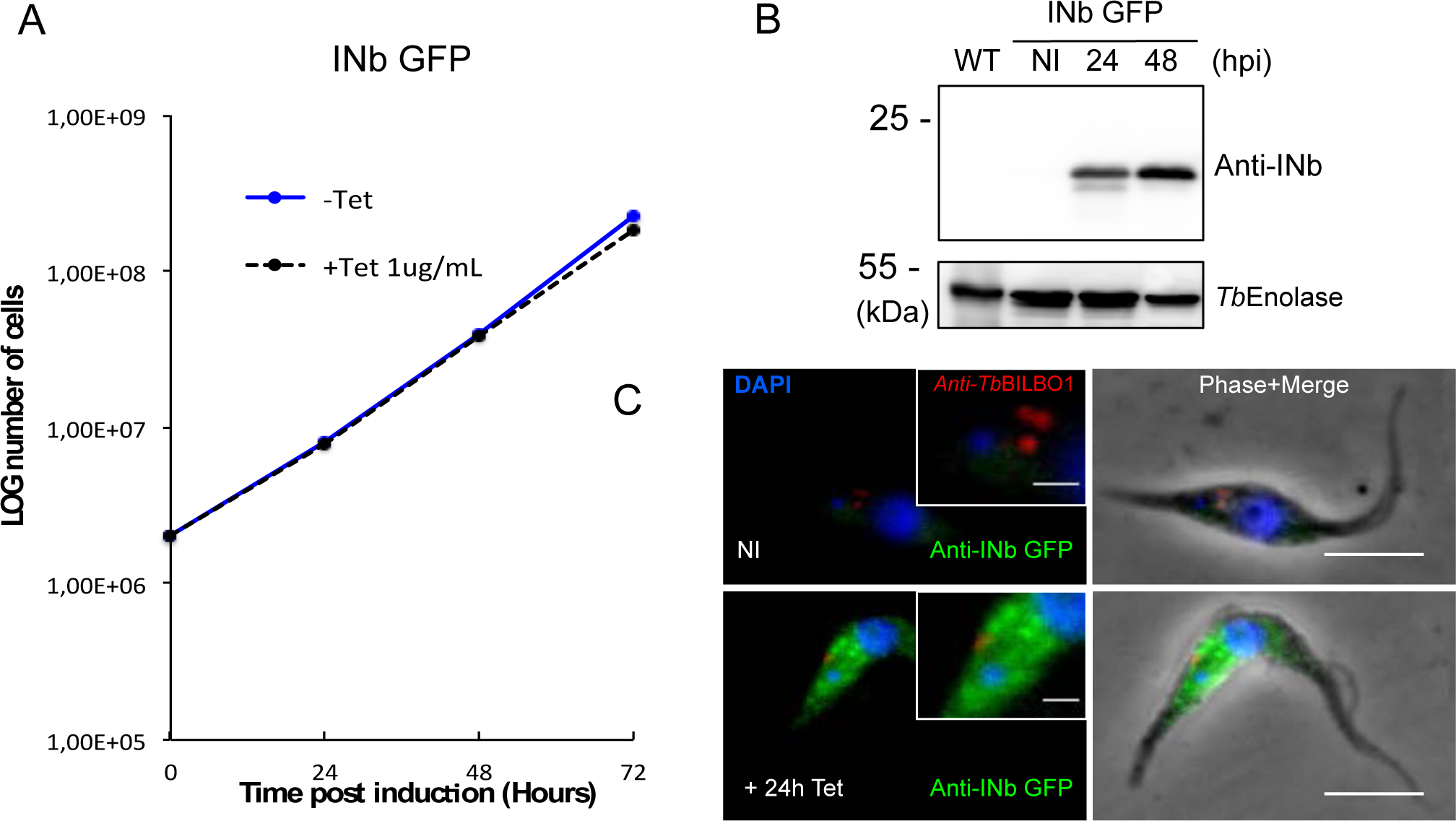
INbGFP::3cMyc expression is cytoplasmic and not lethal to trypanosomes. (A) Growth curve of *T. brucei* PCF cells expressing INbGFP::3cMyc, show that expression of anti-GFP intra-nanobody does not induce a cell growth phenotype. (B) Western blot of expression of INbGFP::3cMyc in *T. brucei* showing expression at 24 and 48hpi. *Tb*Enolase, is a cytoplasmic protein, that has been used as a loading control. (C) IFA of non-induced PCF *T. brucei* cells probed with anti-*Tb*BILBO1 (red) and anti-cMyc to label INbGFP::3cMyc (green - very weak background signal) (-Tet; top panel). A trypanosome induced for 24h (+Tet) demonstrating cytoplasmic localisation of IN-bGFP::3cMyc (green) and the normal FPC localisation of *Tb*BILBO1 (red) (Bottom panel). Scale bar = 5μm. For the sake of convenience, the protein being probed is named as anti-Nb, plus the relevant nanobody name rather than its tag.

**Sup Fig 3.**
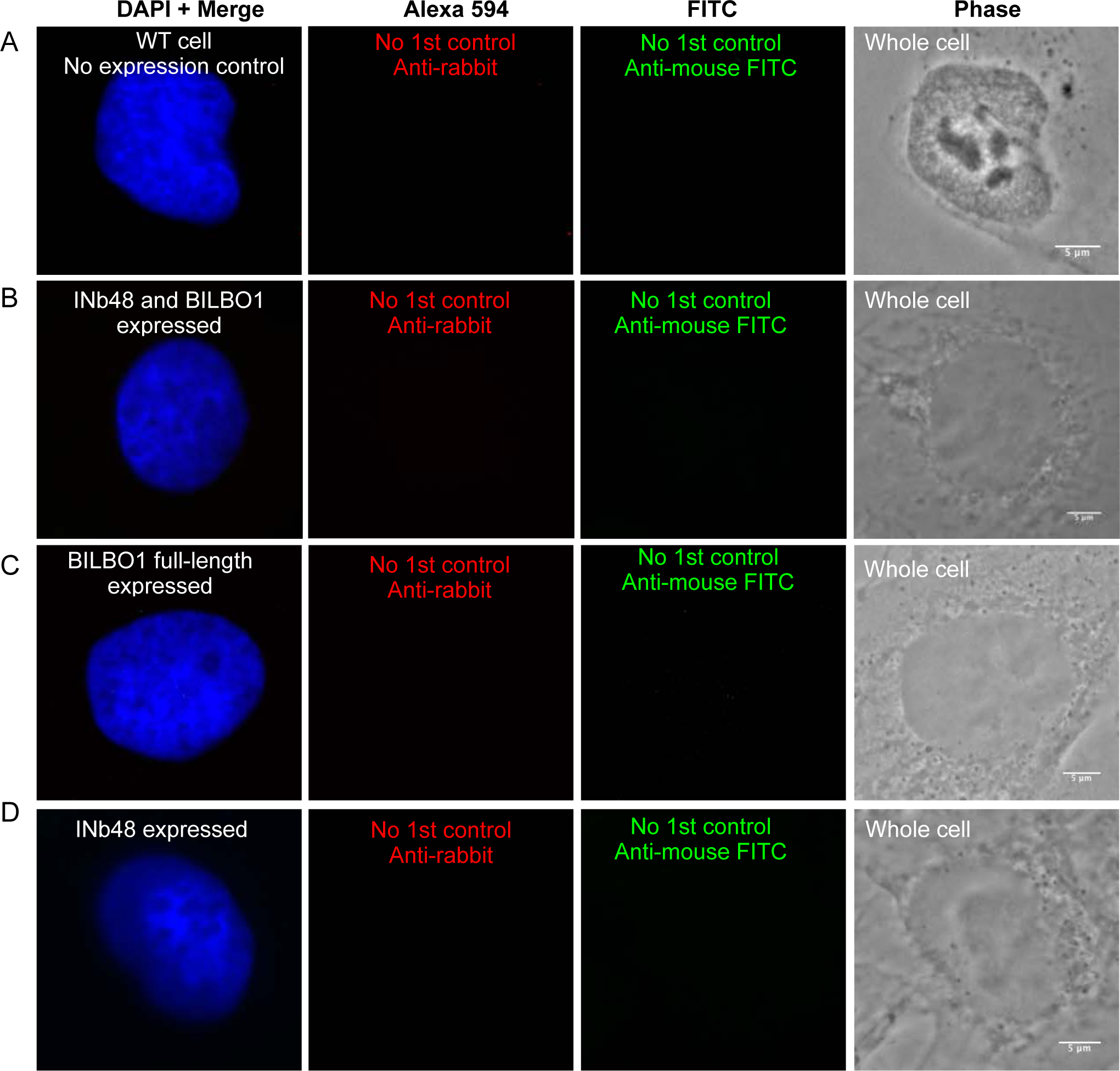
IFA antibody controls for expression of INb48::3HA and/or *Tb*BILBO1 in U-2 OS cells. Controls for Figures 9 and 10. (A) Images of a U-2 OS wild-type (WT) whole cell that has not been transfected with either INb48::3HA or *Tb*BILBO1 and is probed with secondary antibodies only. No signal is observed. (B) Images of a U-2 OS whole cell that has been transfected with INb48::3HA and *Tb*BILBO1 and probed as in (A); no signal is observed. (C) Images of a U-2 OS whole cell that has been transfected with *Tb*BILBO1 full-length and probed as in (A); no signal is observed. (D) Images of a U-2 OS whole cell that has been transfected with INb48::3HA and probed as in (A); no signal is observed. Scale bars = 5μm.

**Sup Fig 4.**
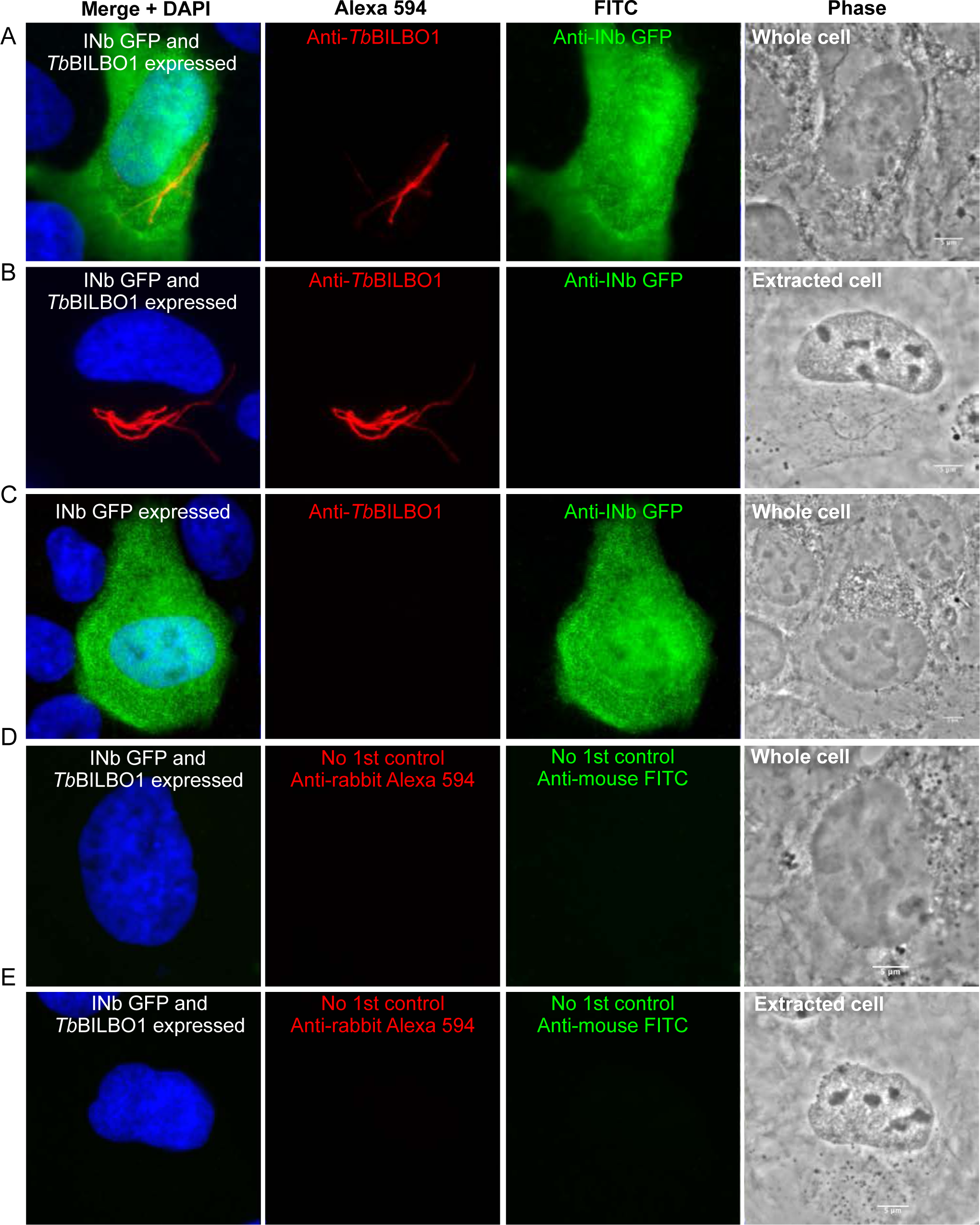
Controls of intra-nanobody anti-GFP (INbGFP**::3HA**) and/or *Tb*BILBO1 expressed in U-2 OS cells. (A) Images of antibody controls of intrabody anti-GFP (INbGFP::3HA) and *Tb*BILBO1 expressed in U-2 OS cells and probed with anti-HA, plus anti-*Tb*BILBO1 (1–110) followed by their respective secondary antibodies anti-rabbit Alexa 594 and anti-mouse FITC. Both *Tb*BILBO1 polymers and the cytoplasmic nanobody (INbGFP::3HA ; green) are observed, but the GFP intrabody does not bind to *Tb*BILBO1 positive polymers (red). (B) Images of a cell treated and probed as in (A) but detergent extracted, illustrating that the anti-GFP intrabody is cytoplasmic and soluble and is removed by detergent extraction but not the *Tb*BILBO1 polymers (red). (C) Images of a U-2 OS cell only expressing the INbGFP::3HA but probed as in (A) and illustrating a green cytoplasmic labelling of the intrabody only, which does not form polymers. (D and E) Images of antibody controls of INbGFP::3HA and *Tb*BILBO1 expressed in U-2 OS cells and probed with secondary antibodies only: Alexa 594 and anti-mouse FITC. (D) is a whole cell whilst (E) is detergent extracted. No signal is observed in either cell. Scale bar = 5μm.

## References

1. Diall O, Cecchi G, Wanda G, Argilés-Herrero R, Vreysen MJB, Cattoli G, Viljoen GJ, Mattioli R, Bouyer J. 2017. Developing a Progressive Control Pathway for African Animal Trypanosomosis. Trends Parasitol 33:499–509.

2. Simarro PP, Cecchi G, Franco JR, Paone M, Diarra A, Priotto G, Mattioli RC, Jannin JG. 2015. Monitoring the Progress towards the Elimination of Gambiense Human African Trypanosomiasis. PLoS Negl Trop Dis 9:e0003785.

3. Mesu VKBK, Kalonji WM, Bardonneau C, Mordt OV, Blesson S, Simon F, Delhomme S, Bernhard S, Kuziena W, Lubaki J-PF, Vuvu SL, Ngima PN, Mbembo HM, Ilunga M, Bonama AK, Heradi JA, Solomo JLL, Mandula G, Badibabi LK, Dama FR, Lukula PK, Tete DN, Lumbala C, Scherrer B, Strub-Wourgaft N, Tarral A. 2018. Oral fexinidazole for late-stage African Trypanosoma brucei gambiense trypanosomiasis: a pivotal multicentre, randomised, non-inferiority trial. Lancet 391:144–154.

4. Deeks ED. 2019. Fexinidazole: First Global Approval. Drugs 79:215–220.

5. Keating J, Yukich JO, Sutherland CS, Woods G, Tediosi F. 2015. Human African trypanosomiasis prevention, treatment and control costs: a systematic review. Acta Trop 150:4–13.

6. Scott V, Sherwin T, Gull K. 1997. gamma-tubulin in trypanosomes: molecular characterisation and localisation to multiple and diverse microtubule organising centres. Journal of cell science 110 ( Pt 2):157–68.

7. Bastin P, Sherwin T, Gull K. 1998. Paraflagellar rod is vital for trypanosome motility. Nature 391:548.

8. Bastin P, Pullen TJ, Sherwin T, Gull K. 1999. Protein transport and flagellum assembly dynamics revealed by analysis of the paralysed trypanosome mutant snl-1. Journal of cell science 112 ( Pt 21):3769–77.

9. Kohl L, Sherwin T, Gull K. 1999. Assembly of the paraflagellar rod and the flagellum attachment zone complex during the Trypanosoma brucei cell cycle. J Eukaryot Microbiol 46:105–9.

10. Moreira-Leite FF, Sherwin T, Kohl L, Gull K. 2001. A trypanosome structure involved in transmitting cytoplasmic information during cell division. Science (New York, NY 294:610–2.

11. Robinson DR, Gull K. 1991. Basal body movements as a mechanism for mitochondrial genome segregation in the trypanosome cell cycle. Nature 352:731–3.

12. Vedrenne C, Giroud C, Robinson DR, Besteiro S, Bosc C, Bringaud F, Baltz T. 2002. Two related subpellicular cytoskeleton-associated proteins in Trypanosoma brucei stabilize microtubules. Molecular biology of the cell 13:1058–70.

13. Pradel LC, Bonhivers M, Landrein N, Robinson DR. 2006. NIMA-related kinase TbNRKC is involved in basal body separation in Trypanosoma brucei. Journal of cell science 119:1852–63.

14. Bonhivers M, Nowacki S, Landrein N, Robinson DR. 2008. Biogenesis of the trypanosome endo-exocytotic organelle is cytoskeleton mediated. PLoS Biol 6:e105.

15. Henley GL, Lee CM, Takeuchi A. 1978. Electron microscopy observations on Trypanosoma brucei: freeze-cleaving and thin-sectioning study of the apical part of the flagellar pocket. Z Parasitenkd 55:181–7.

16. Webster P, Russell DG. 1993. The flagellar pocket of trypanosomatids. Parasitology today (Personal ed 9:201–6.

17. Morgan GW, Hall BS, Denny PW, Carrington M, Field MC. 2002. The kinetoplastida endocytic apparatus. Part I: a dynamic system for nutrition and evasion of host defences. Trends in parasitology 18:491–6.

18. Field MC, Carrington M. 2009. The trypanosome flagellar pocket. Nature reviews 7:775–86.

19. Alcantara C de L, Vidal JC, de Souza W, Cunha-E-Silva NL. 2017. The cytostome-cytopharynx complex of Trypanosoma cruzi epimastigotes disassembles during cell division. J Cell Sci 130:164–176.

20. Esson HJ, Morriswood B, Yavuz S, Vidilaseris K, Dong G, Warren G. 2012. Morphology of the trypanosome bilobe, a novel cytoskeletal structure. Eukaryotic cell 11:761–72.

21. Vidilaseris K, Morriswood B, Kontaxis G, Dong G. 2014. Structure of the TbBILBO1 protein N-terminal domain from Trypanosoma brucei reveals an essential requirement for a conserved surface patch. J Biol Chem 289:3724–3735.

22. Vidilaseris K, Shimanovskaya E, Esson HJ, Morriswood B, Dong G. 2014. Assembly mechanism of Trypanosoma brucei BILBO1, a multidomain cytoskeletal protein. J Biol Chem 289:23870–23881.

23. Florimond C, Sahin A, Vidilaseris K, Dong G, Landrein N, Dacheux D, Albisetti A, Byard EH, Bonhivers M, Robinson DR. 2015. BILBO1 is a scaffold protein of the flagellar pocket collar in the pathogen Trypanosoma brucei. PLoS Pathog 11:e1004654.

24. Perdomo D, Bonhivers M, Robinson DR. 2016. The Trypanosome Flagellar Pocket Collar and Its Ring Forming Protein-TbBILBO1. Cells 5.

25. Albisetti A, Florimond C, Landrein N, Vidilaseris K, Eggenspieler M, Lesigang J, Dong G, Robinson DR, Bonhivers M. 2017. Interaction between the flagellar pocket collar and the hook complex via a novel microtubule-binding protein in Trypanosoma brucei. PLoS Pathog 13:e1006710.

26. Vidilaseris K, Landrein N, Pivovarova Y, Lesigang J, Aeksiri N, Robinson DR, Bonhivers M, Dong G. 2020. Crystal structure of the N-terminal domain of the trypanosome flagellar protein BILBO1 reveals a ubiquitin fold with a long structured loop for protein binding. J Biol Chem 295:1489–1499.

27. Morriswood B, He CY, Sealey-Cardona M, Yelinek J, Pypaert M, Warren G. 2009. The bilobe structure of Trypanosoma brucei contains a MORN-repeat protein. Molecular and biochemical parasitology 167:95–103.

28. Morriswood B, Havlicek K, Demmel L, Yavuz S, Sealey-Cardona M, Vidilaseris K, Anrather D, Kostan J, Djinovic-Carugo K, Roux KJ, Warren G. 2013. Novel Bilobe Components in Trypanosoma brucei Identified Using Proximity-Dependent Biotinylation. Eukaryotic cell 12:356–67.

29. Morriswood B. 2015. Form, Fabric, and Function of a Flagellum-Associated Cytoskeletal Structure. Cells 4:726–747.

30. Morriswood B, Schmidt K. 2015. A MORN Repeat Protein Facilitates Protein Entry into the Flagellar Pocket of Trypanosoma brucei. Eukaryotic Cell 14:1081–1093.

31. Hamers-Casterman C, Atarhouch T, Muyldermans S, Robinson G, Hamers C, Songa EB, Bendahman N, Hamers R. 1993. Naturally occurring antibodies devoid of light chains. Nature 363:446–448.

32. Greenberg AS, Avila D, Hughes M, Hughes A, McKinney EC, Flajnik MF. 1995. A new antigen receptor gene family that undergoes rearrangement and extensive somatic diversification in sharks. Nature 374:168–173.

33. Nguyen VK, Desmyter A, Muyldermans S. 2001. Functional heavy-chain antibodies in Camelidae. Adv Immunol 79:261–296.

34. Holliger P, Hudson PJ. 2005. Engineered antibody fragments and the rise of single domains. Nat Biotechnol 23:1126–1136.

35. Ward ES, Güssow D, Griffiths AD, Jones PT, Winter G. 1989. Binding activities of a repertoire of single immunoglobulin variable domains secreted from Escherichia coli. Nature 341:544–546.

36. Muyldermans S. 2013. Nanobodies: natural single-domain antibodies. Annu Rev Biochem 82:775–797.

37. Mir MA, Mehraj U, Sheikh BA, Hamdani SS. 2019. Nanobodies: The “magic bullets” in therapeutics, drug delivery and diagnostics. Hum Antibodies https://doi.org/10.3233/HAB-190390.

38. Matz H, Dooley H. 2019. Shark IgNAR-derived binding domains as potential diagnostic and therapeutic agents. Dev Comp Immunol 90:100–107.

39. Woods J. 2019. Selection of Functional Intracellular Nanobodies. SLAS Discov 24:703–713.

40. Arias JL, Unciti-Broceta JD, Maceira J, Del Castillo T, Hernández-Quero J, Magez S, Soriano M, García-Salcedo JA. 2015. Nanobody conjugated PLGA nanoparticles for active targeting of African Trypanosomiasis. J Control Release 197:190–198.

41. Pardon E, Laeremans T, Triest S, Rasmussen SGF, Wohlkönig A, Ruf A, Muyldermans S, Hol WGJ, Kobilka BK, Steyaert J. 2014. A general protocol for the generation of Nanobodies for structural biology. Nat Protoc 9:674–693.

42. Van Audenhove I, Van Impe K, Ruano-Gallego D, De Clercq S, De Muynck K, Vanloo B, Verstraete H, Fernández LÁ, Gettemans J. 2013. Mapping cytoskeletal protein function in cells by means of nanobodies. Cytoskeleton (Hoboken) 70:604–622.

43. Muyldermans S, Baral TN, Retamozzo VC, De Baetselier P, De Genst E, Kinne J, Leonhardt H, Magez S, Nguyen VK, Revets H, Rothbauer U, Stijlemans B, Tillib S, Wernery U, Wyns L, Hassanzadeh-Ghassabeh G, Saerens D. 2009. Camelid immunoglobulins and nanobody technology. Vet Immunol Immunopathol 128:178–83.

44. Abderrazek RB, Hmila I, Vincke C, Benlasfar Z, Pellis M, Dabbek H, Saerens D, El Ayeb M, Muyldermans S, Bouhaouala-Zahar B. 2009. Identification of potent nanobodies to neutralize the most poisonous polypeptide from scorpion venom. The Biochemical journal 424:263–72.

45. Kaplon H, Reichert JM. 2019. Antibodies to watch in 2019. MAbs 11:219–238.

46. Duggan S. 2018. Caplacizumab: First Global Approval. Drugs 78:1639–1642.

47. Morrison C. 2019. Nanobody approval gives domain antibodies a boost. Nat Rev Drug Discov 18:485–487.

48. Cardinale A, Filesi I, Vetrugno V, Pocchiari M, Sy M-S, Biocca S. 2005. Trapping prion protein in the endoplasmic reticulum impairs PrPC maturation and prevents PrPSc accumulation. J Biol Chem 280:685–694.

49. Marschall ALJ, Dübel S. 2016. Antibodies inside of a cell can change its outside: Can intrabodies provide a new therapeutic paradigm? Comput Struct Biotechnol J 14:304–308.

50. Messer A, Joshi SN. 2013. Intrabodies as neuroprotective therapeutics. Neurotherapeutics 10:447–458.

51. Alzogaray V, Danquah W, Aguirre A, Urrutia M, Berguer P, Garcia Vescovi E, Haag F, Koch-Nolte F, Goldbaum FA. 2010. Single-domain llama antibodies as specific intracellular inhibitors of SpvB, the actin ADP-ribosylating toxin of Salmonella typhimurium. FASEB J https://doi.org/fj.10-162958 [pii] 10.1096/fj.10-162958.

52. Zhou C, Przedborski S. 2009. Intrabody and Parkinson’s disease. Biochimica et biophysica acta 1792:634–42.

53. Pinto Torres JE, Goossens J, Ding J, Li Z, Lu S, Vertommen D, Naniima P, Chen R, Muyldermans S, Sterckx YG-J, Magez S. 2018. Development of a Nanobody-based lateral flow assay to detect active Trypanosoma congolense infections. Sci Rep 8:9019.

54. Van Impe K, Bethuyne J, Cool S, Impens F, Ruano-Gallego D, De Wever O, Vanloo B, Van Troys M, Lambein K, Boucherie C, Martens E, Zwaenepoel O, Hassanzadeh-Ghassabeh G, Vandekerckhove J, Gevaert K, Fernández LÁ, Sanders NN, Gettemans J. 2013. A nanobody targeting the F-actin capping protein CapG restrains breast cancer metastasis. Breast Cancer Res 15:R116.

55. Meli G, Krako N, Manca A, Lecci A, Cattaneo A. 2013. Intrabodies for protein interference in Alzheimer’s disease. J Biol Regul Homeost Agents 27:89–105.

56. Alirahimi E, Ashkiyan A, Kazemi-Lomedasht F, Azadmanesh K, Hosseininejad-Chafi M, Habibi-Anbouhi M, Moazami R, Behdani M. 2017. Intrabody targeting vascular endothelial growth factor receptor-2 mediates downregulation of surface localization. Cancer Gene Ther 24:33–37.

57. Aires da Silva F, Santa-Marta M, Freitas-Vieira A, Mascarenhas P, Barahona I, Moniz-Pereira J, Gabuzda D, Goncalves J. 2004. Camelized rabbit-derived VH single-domain intrabodies against Vif strongly neutralize HIV-1 infectivity. Journal of molecular biology 340:525–42.

58. Reimer E, Somplatzki S, Zegenhagen D, Hänel S, Fels A, Bollhorst T, Hovest LG, Bauer S, Kirschning CJ, Böldicke T. 2013. Molecular cloning and characterization of a novel anti-TLR9 intrabody. Cell Mol Biol Lett 18:433–446.

59. Stijlemans B, Conrath K, Cortez-Retamozo V, Van Xong H, Wyns L, Senter P, Revets H, De Baetselier P, Muyldermans S, Magez S. 2004. Efficient targeting of conserved cryptic epitopes of infectious agents by single domain antibodies. African trypanosomes as paradigm. J Biol Chem 279:1256–1261.

60. Baral TN, Magez S, Stijlemans B, Conrath K, Vanhollebeke B, Pays E, Muyldermans S, De Baetselier P. 2006. Experimental therapy of African trypanosomiasis with a nanobody-conjugated human trypanolytic factor. Nat Med 12:580–584.

61. Saerens D, Stijlemans B, Baral TN, Nguyen Thi GT, Wernery U, Magez S, De Baetselier P, Muyldermans S, Conrath K. 2008. Parallel selection of multiple anti-infectome Nanobodies without access to purified antigens. J Immunol Methods 329:138–50.

62. Stijlemans B, Caljon G, Natesan SKA, Saerens D, Conrath K, Pérez-Morga D, Skepper JN, Nikolaou A, Brys L, Pays E, Magez S, Field MC, De Baetselier P, Muyldermans S. 2011. High affinity nanobodies against the Trypanosome brucei VSG are potent trypanolytic agents that block endocytosis. PLoS Pathog 7:e1002072.

63. Caljon G, Caveliers V, Lahoutte T, Stijlemans B, Ghassabeh GH, Van Den Abbeele J, Smolders I, De Baetselier P, Michotte Y, Muyldermans S, Magez S, Clinckers R. 2012. Using microdialysis to analyse the passage of monovalent nanobodies through the blood-brain barrier. Br J Pharmacol 165:2341–2353.

64. Caljon G, Stijlemans B, Saerens D, Van Den Abbeele J, Muyldermans S, Magez S, De Baetselier P. 2012. Affinity is an important determinant of the anti-trypanosome activity of nanobodies. PLoS Negl Trop Dis, 2012/11/15 ed. 6:e1902–e1902.

65. Ditlev SB, Florea R, Nielsen MA, Theander TG, Magez S, Boeuf P, Salanti A. 2014. Utilizing nanobody technology to target non-immunodominant domains of VAR2CSA. PLoS One 9:e84981–e84981.

66. Obishakin E, Stijlemans B, Santi-Rocca J, Vandenberghe I, Devreese B, Muldermans S, Bastin P, Magez S. 2014. Generation of a nanobody targeting the paraflagellar rod protein of trypanosomes. PLoS ONE 9:e115893.

67. Unciti-Broceta JD, Arias JL, Maceira J, Soriano M, Ortiz-González M, Hernández-Quero J, Muñóz-Torres M, de Koning HP, Magez S, Garcia-Salcedo JA. 2015. Specific Cell Targeting Therapy Bypasses Drug Resistance Mechanisms in African Trypanosomiasis. PLoS Pathog 11:e1004942.

68. Odongo S, Sterckx YGJ, Stijlemans B, Pillay D, Baltz T, Muyldermans S, Magez S. 2016. An Anti-proteome Nanobody Library Approach Yields a Specific Immunoassay for Trypanosoma congolense Diagnosis Targeting Glycosomal Aldolase. PLoS Negl Trop Dis 10:e0004420.

69. Stijlemans B, De Baetselier P, Caljon G, Van Den Abbeele J, Van Ginderachter JA, Magez S. 2017. Nanobodies As Tools to Understand, Diagnose, and Treat African Trypanosomiasis. Front Immunol 8:724.

70. Vickerman K. 1969. The fine structure of Trypanosoma congolense in its bloodstream phase. J Protozool 16:54–69.

71. Dong X, Lim TK, Lin Q, He CY. 2020. Basal Body Protein TbSAF1 Is Required for Microtubule Quartet Anchorage to the Basal Bodies in Trypanosoma brucei. mBio 11:e00668–20.

72. Gheiratmand L, Brasseur A, Zhou Q, He CY. 2013. Biochemical Characterization of the Bilobe Reveals a Continuous Structural Network Linking the Bi-lobe to Other Single-copied Organelles in Trypanosoma brucei. The Journal of biological chemistry 288:3489–99.

73. Wheeler RJ, Gull K, Sunter JD. 2019. Coordination of the Cell Cycle in Trypanosomes. Annu Rev Microbiol 73:133–154.

74. Bastin P, MacRae TH, Francis SB, Matthews KR, Gull K. 1999. Flagellar morphogenesis: protein targeting and assembly in the paraflagellar rod of trypanosomes. Molecular and cellular biology 19:8191–200.

75. Dang HQ, Zhou Q, Rowlett VW, Hu H, Lee KJ, Margolin W, Li Z. 2017. Proximity Interactions among Basal Body Components in Trypanosoma brucei Identify Novel Regulators of Basal Body Biogenesis and Inheritance. mBio 8.

76. Van den Abbeele A, De Clercq S, De Ganck A, De Corte V, Van Loo B, Soror SH, Srinivasan V, Steyaert J, Vandekerckhove J, Gettemans J. 2010. A llama-derived gelsolin single-domain antibody blocks gelsolin-G-actin interaction. Cell Mol Life Sci 67:1519–1535.

77. De Vooght L, Caljon G, Stijlemans B, De Baetselier P, Coosemans M, Van den Abbeele J. 2012. Expression and extracellular release of a functional anti-trypanosome Nanobody® in Sodalis glossinidius, a bacterial symbiont of the tsetse fly. Microb Cell Fact 11:23.

78. De Vooght L, Caljon G, De Ridder K, Van Den Abbeele J. 2014. Delivery of a functional anti-trypanosome Nanobody in different tsetse fly tissues via a bacterial symbiont, Sodalis glossinidius. Microb Cell Fact 13:156–156.

79. Böldicke T. 2017. Single domain antibodies for the knockdown of cytosolic and nuclear proteins. Protein Sci, 2017/03/24 ed. 26:925–945.

80. Koo MY, Park J, Lim JM, Joo SY, Shin S-P, Shim HB, Chung J, Kang D, Woo HA, Rhee SG. 2014. Selective inhibition of the function of tyrosine-phosphorylated STAT3 with a phosphorylation site-specific intrabody. Proc Natl Acad Sci U S A, 2014/04/14 ed. 111:6269–6274.

81. Stanimirovic D, Kemmerich K, Haqqani AS, Farrington GK. 2014. Engineering and pharmacology of blood-brain barrier-permeable bispecific antibodies. Adv Pharmacol 71:301–335.

82. Abulrob A, Sprong H, Van Bergen en Henegouwen P, Stanimirovic D. 2005. The blood-brain barrier transmigrating single domain antibody: mechanisms of transport and antigenic epitopes in human brain endothelial cells. J Neurochem 95:1201–1214.

83. Negi SS, Braun W. 2017. Cross-React: a new structural bioinformatics method for predicting allergen cross-reactivity. Bioinformatics 33:1014–1020.

84. Faktorová D, Nisbet RER, Fernández Robledo JA, Casacuberta E, Sudek L, Allen AE, Ares M Jr, Aresté C, Balestreri C, Barbrook AC, Beardslee P, Bender S, Booth DS, Bouget F-Y, Bowler C, Breglia SA, Brownlee C, Burger G, Cerutti H, Cesaroni R, Chiurillo MA, Clemente T, Coles DB, Collier JL, Cooney EC, Coyne K, Docampo R, Dupont CL, Edgcomb V, Einarsson E, Elustondo PA, Federici F, Freire-Beneitez V, Freyria NJ, Fukuda K, García PA, Girguis PR, Gomaa F, Gornik SG, Guo J, Hampl V, Hanawa Y, Haro-Contreras ER, Hehenberger E, High-field A, Hirakawa Y, Hopes A, Howe CJ, Hu I, Ibañez J, Irwin NAT, Ishii Y, Janowicz NE, Jones AC, Kachale A, Fujimura-Kamada K, Kaur B, Kaye JZ, Kazana E, Keeling PJ, King N, Klobutcher LA, Lander N, Lassadi I, Li Z, Lin S, Lozano J-C, Luan F, Maruyama S, Matute T, Miceli C, Minagawa J, Moosburner M, Najle SR, Nanjappa D, Nimmo IC, Noble L, Novák Vanclová AMG, Nowacki M, Nuñez I, Pain A, Piersanti A, Pucciarelli S, Pyrih J, Rest JS, Rius M, Robertson D, Ruaud A, Ruiz-Trillo I, Sigg MA, Silver PA, Slamovits CH, Jason Smith G, Sprecher BN, Stern R, Swart EC, Tsaousis AD, Tsypin L, Turkewitz A, Turnšek J, Valach M, Vergé V, von Dassow P, von der Haar T, Waller RF, Wang L, Wen X, Wheeler G, Woods A, Zhang H, Mock T, Worden AZ, Lukeš J. 2020. Genetic tool development in marine protists: emerging model organisms for experimental cell biology. Nat Methods, 2020/04/06 ed. 17:481–494.

85. Wirtz E, Leal S, Ochatt C, Cross GA. 1999. A tightly regulated inducible expression system for conditional gene knock-outs and dominant-negative genetics in Trypanosoma brucei. Molecular and biochemical parasitology 99:89–101.

86. Nossal NG, Heppel LA. 1966. The release of enzymes by osmotic shock from Escherichia coli in exponential phase. J Biol Chem 241:3055–3062.

87. Karlsson R, Katsamba PS, Nordin H, Pol E, Myszka DG. 2006. Analyzing a kinetic titration series using affinity biosensors. Anal Biochem 349:136–147.

88. Palau W, Di Primo C. 2012. Single-cycle kinetic analysis of ternary DNA complexes by surface plasmon resonance on a decaying surface. Biochimie 94:1891–1899.

89. Schumann Burkard G, Jutzi P, Roditi I. 2011. Genome-wide RNAi screens in bloodstream form trypanosomes identify drug transporters. Mol Biochem Parasitol 175:91–94.

90. Heldin CH, Johnsson A, Wennergren S, Wernstedt C, Betsholtz C, Westermark B. 1986. A human osteosarcoma cell line secretes a growth factor structurally related to a homodimer of PDGF A-chains. Nature 319:511–4.

91. Dacheux D, Landrein N, Thonnus M, Gilbert G, Sahin A, Wodrich H, Robinson DR, Bonhivers M. 2012. A MAP6-Related Protein Is Present in Protozoa and Is Involved in Flagellum Motility. PloS one 7:e31344.

92. Hannaert V, Albert M-A, Rigden DJ, da Silva Giotto MT, Thiemann O, Garratt RC, Van Roy J, Opperdoes FR, Michels PAM. 2003. Kinetic characterization, structure modelling studies and crystallization of Trypanosoma brucei enolase. Eur J Biochem 270:3205–3213.

93. Woods A, Sherwin T, Sasse R, MacRae TH, Baines AJ, Gull K. 1989. Definition of individual components within the cytoskeleton of Trypanosoma brucei by a library of monoclonal antibodies. Journal of cell science 93 ( Pt 3):491–500.

94. Poon SK, Peacock L, Gibson W, Gull K, Kelly S. 2012. A modular and optimized single marker system for generating Trypanosoma brucei cell lines expressing T7 RNA polymerase and the tetracycline repressor. Open Biol 2:110037–110037.

